# Developmentally-regulated proteolysis by MdfA and ClpCP mediates metabolic differentiation during *Bacillus subtilis* sporulation

**DOI:** 10.1101/2024.11.26.625531

**Authors:** Eammon P. Riley, Jelani A. Lyda, Octavio Reyes-Matte, Joseph Sugie, Iqra R. Kasu, Eray Enustun, Emily Armbruster, Sumedha Ravishankar, Rivka L. Isaacson, Amy H. Camp, Javier Lopez-Garrido, Kit Pogliano

**Author notes:** These authors contributed equally to this work (Eammon P. Riley, Jelani A. Lyda). Correspondence to Kit Pogliano or Javier Lopez Garrido.

## Abstract

*Bacillus subtilis* sporulation entails a dramatic transformation of the two cells required to assemble a dormant spore, with the larger mother cell engulfing the smaller forespore to produce the cell-within-a-cell structure that is a hallmark of endospore formation. Sporulation also entails metabolic differentiation, whereby key metabolic enzymes are depleted from the forespore but maintained in the mother cell. This reduces the metabolic potential of the forespore, which becomes dependent on mother-cell metabolism and the SpoIIQ-SpoIIIA channel to obtain metabolic building blocks necessary for development. We demonstrate that metabolic differentiation depends on the ClpCP protease and a forespore-produced protein encoded by the *yjbA* gene, which we have renamed MdfA (metabolic differentiation factor A). MdfA is conserved in aerobic endospore-formers and required for spore resistance to hypochlorite. Using mass spectrometry and quantitative fluorescence microscopy, we show that MdfA mediates the depletion of dozens of metabolic enzymes and key transcription factors from the forespore. An accompanying study by Massoni, Evans and collaborators demonstrates that MdfA is a ClpC adaptor protein that directly interacts with and stimulates ClpCP activity. Together, these results document a developmentally-regulated proteolytic pathway that reshapes forespore metabolism, reinforces differentiation, and is required to produce spores resistant to the oxidant hypochlorite.

## Introduction

The endospore formation pathway of *Bacillus subtilis* and its relatives is a striking example of how bacteria can deploy their streamlined cellular machineries to mediate dramatic morphological transformations (Tan and Ramamurthi 2014; Khanna et al. 2020; Riley et al. 2021b). Like many eukaryotic developmental pathways, endospore formation commences with an asymmetric cell division (Levin and Losick 1996; Errington 2003; Barák et al. 2019) that produces two cells of differing size and fate: the smaller forespore that will become a dormant spore, and the larger mother cell that lyses after contributing to spore development (Fig. 1A). The early forespore- and mother cell-specific transcription factors s^F^ and s^E^ are activated shortly after polar septation (Piggot and Losick 2001; Hilbert and Piggot 2004), and the two cells initiate a phagocytosis-like process known as engulfment that is a hallmarks of endospore formation. Engulfment is mediated by coordinated, cell-specific peptidoglycan degradation and synthesis that remodels the cell wall and allows the mother cell membrane to migrate around the forespore until the future spore is completely enclosed in the mother cell cytoplasm (Abanes-De Mello et al. 2002; Gutierrez et al. 2010; Morlot et al. 2010; Tocheva et al. 2013; Ojkic et al. 2016; Khanna et al. 2019; Chan et al. 2022). The late forespore- and mother cell-specific transcription factors s^G^ and s^K^ are then activated to direct synthesis of proteins that complete assembly of the spore envelope, protect the spore chromosome from UV damage, dehydrate the spore cytoplasm, and complete the transition of the spore to dormancy (Popham and Bernhards 2015; Driks and Eichenberger 2016; Setlow and Christie 2023). Ultimately, the mother cell lyses to release the mature spore into the environment, where it remains dormant until nutrients become available, when it can rapidly resume growth (Moir and Cooper 2015; Christie and Setlow 2020).

**Figure 1.**
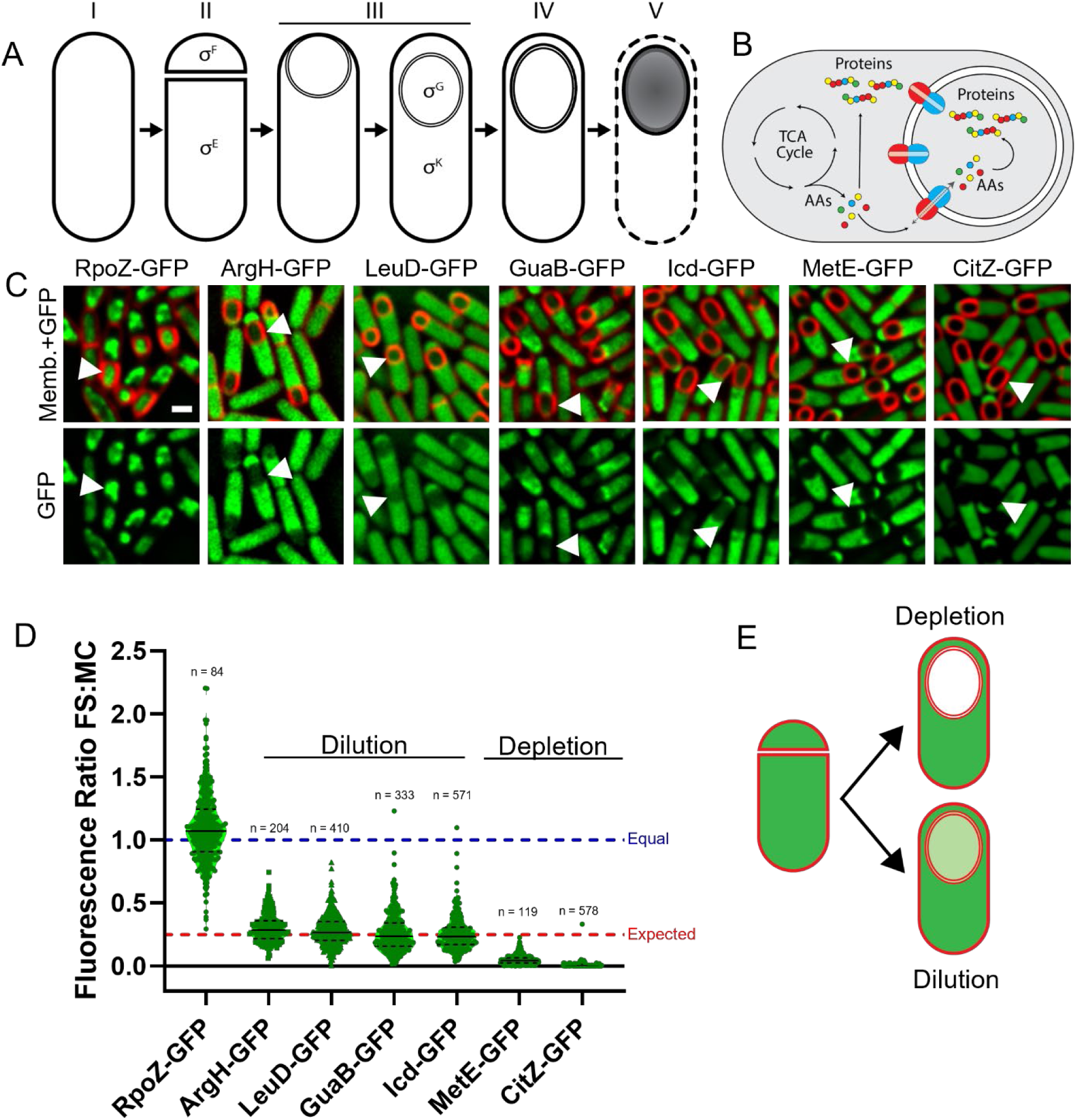
Metabolic reprogramming during spore formation. (A) *B. subtills* sporulation. I, vegetative cell; II, polar septation; III, engulfment; IV, spore maturation; V, mother cell lysis and spore release. Sporulation-specific sigma (σ) are shown in the cells and stages they are active. (B) Intercellular nurturing during *B. subtilis* sporulation. Mother cell TCA cycle and amino acid synthesis (AAs, colored circles) enables forespore protein synthesis. Metabolites are transported via the Q-A channel (blue, SpoIIQ: red, SpoIIIAA-AH). (C) Fluorescence micrographs of sporulating cells with FM 4-64 stained membranes (red) and GFP fusions (green) to proteins involved in transcription (RpoZ), TCA cycle (Icd and CitZ), nucleotide biosynthesis (GuaB) and amino acid biosynthesis (ArgH, LeuD and MetE). Arrowheads indicate representative forespores. Scale bar, 1 µm. (D) Violin plots showing the ratios of the mean GFP fluorescence in the forespore to that in the mother cell (FS:MC) for the proteins in (C). Each green dot represents the fluorescent ratio of an individual sporangium. The black dotted lines delimit the middle 75% of the data and the black solid lines indicate the median. The blue dashed line indicates equal intensity in both cells, the red dashed line indicates a fluorescence ratio of 0.25, as predicted for protein dilution due to increased forespore volume and decreased mother cell volume (Fig. S1). Values below 0.25 likely entail protein depletion. (E) Diagram showing the two pathways, dilution due to forespore growth, and a specific protein depletion pathway.

Sporulation therefore entails the dual challenges of transitioning one cell to metabolic dormancy while synthesizing large quantities of proteins in both cells that are needed to assemble a durable spore, and maintaining the ability of the spore to rapidly exit dormancy and resume full metabolic activity. Although classical biochemical studies and modern proteomics experiments have shown that spores have significantly lower levels of many metabolic enzymes than growing cells (Kornberg et al. 1968; Swarge et al. 2020), it has remained unclear when during development the spore transitions to dormancy. Given the need for substantial protein synthesis in the forespore from highly expressed s^G^-dependent genes that encode the a/b SASP proteins that render the spore chromosome resistant to UV damage (Setlow 1988; Nerber and Sorg 2024), it has seemed likely that the transition to metabolic dormancy was one of the last steps of sporulation. Contrary to this notion, we demonstrated that the forespore and mother cell become metabolically differentiated shortly after polar septation, when the activation of the early forespore transcription factor s^F^ leads to the rapid depletion of key metabolic enzymes from the forespore (Riley et al. 2021a). This reduces the metabolic potential of the forespore and makes forespore protein synthesis dependent on mother cell metabolic enzymes, including those in the TCA cycle and precursor biosynthesis (Fig. 1B). We showed that amino acids from the mother cell are incorporated into forespore proteins (Riley et al. 2021a), confirming that the mother cell nurtures the forespore, as implied by its name. Thus, *B. subtilis* sporulation entails both the morphological differentiation of the two cells required to make a spore and the metabolic remodeling of the forespore, which depends on and is nurtured by the mother cell during development.

Several lines of evidence suggest that the mother cell nurtures the forespore via the SpoIIQ-SpoIIIA (Q-A) protein complex, which spans the forespore and mother cell membranes and is essential for sporulation (Blaylock et al. 2004; Doan et al. 2009). Camp and Losick showed that the Q-A complex is required for sustained activity of s^F^ and T7 RNA polymerase in the forespore (Camp and Losick 2009), leading the authors to propose that the complex assembles a gap junction-like channel through which the mother cell provides the forespores with metabolites required for gene expression. This hypothesis has been supported by structural data showing that components of the Q-A complex can assemble into a channel (Levdikov et al. 2012; Meisner et al. 2012; Rodrigues et al. 2016; Zeytuni et al. 2017), the accessiblity of mother cell components of the complex to biotinylation from the forespore cytoplasm (Meisner et al. 2008), and our more recent observations that the Q-A complex is required for forespore protein synthesis and for the bidirectional diffusion of the fluorophore calcein between the mother cell and the forespore (Riley et al. 2021a). Altogether, the data support the hypothesis that the Q-A complex is a channel that allows the mother cell to nurture the forespore during sporulation.

The mechanism, extent and impact of metabolic differentiation remains unclear, although timelapse microscopy suggested that at least one protein (CitZ) was actively depleted from the forespore, likely via proteolysis (Riley et al. 2021a). We here demonstrate that two pathways reduce protein concentration in the forespore, a non-specific pathway in which the increased forespore volume dilutes a wide array of proteins, and a specific pathway in which certain proteins are reduced to undetectable levels by a forespore-specific proteolytic event that depends on the ClpCP protease and a forespore-specific protein (YjbA) that we have renamed MdfA for <u>m</u>etabolic <u>d</u>ifferentiation factor A. Mass spectrometry demonstrated that dozens of metabolic enzymes and transcription factors are depleted from spores in an MdfA-dependent manner. We also found that MdfA is required to produce spores that are resistant to NaOCl-induced oxidative stress. Together, our results describe a molecular mechanism that contributes to the metabolic differentiation of the forespore, a new aspect of *B. subtilis* sporulation that is required to produce spores resistant to NaOCl-induced oxidative stress.

## Results

### Two pathways reduce protein concentration in the forespore

We previously showed that key metabolic enzymes are depleted from the forespore after asymmetric division (Riley et al. 2021a). To better understand the mechanisms for the reduction of metabolic potential in the forespore, we performed quantitative fluorescence microscopy on a panel of GFP fusions to different metabolic proteins. We noted striking differences in the extent to which the levels of certain enzymes were reduced in the forespore compared to the mother cell (Fig. 1C and D, (Riley et al. 2021a)), with many proteins such as isocitrate dehydrogenase-GFP (Icd-GFP) decreasing ∼4-fold in the forespore relative to the mother cell (Fig 1D; Riley et al. 2021a), and others such as citrate synthase-GFP (CitZ-GFP) decreasing to undetectable levels (Fig. 1C-D). This suggests that there may be two pathways to decrease protein levels in the forespore (Fig. 1E): (1) a general pathway that yields a widespread ∼4-fold reduction in protein concentration in the forespore compared to the mother cell, and (2) a specific pathway that mediates a more dramatic reduction in the concentration of specific enzymes, such as CitZ, in the forespore.

### Cell growth mediates a general reduction in forespore protein concentration

The general pathway impacted numerous proteins including Icd, LeuD, GuaB and ArgH (Fig. 1D), as well as other proteins in central metabolism and amino acid and nucleotide synthesis (Riley et al. 2021a). This 4-fold reduction in protein concentration in the forespore could be due to the fact that the forespore grows during sporulation, roughly tripling its initial volume, while the volume of the mother cell decreases slightly due to forespore growth (Lopez-Garrido et al. 2018). In the absence of new protein synthesis, this would cause proteins to be diluted in the forespore and concentrated in the mother cell, resulting in a forespore-to-mother cell (FS:MC) protein concentration ratio of ∼0.25 (Fig. S1). Resynthesis of proteins in the forespore could increase this ratio, as illustrated by RpoZ-GFP, whose production is boosted in the forespore by σ^F^ (Abecasis et al. 2013), and shows a similar concentration in the two cells (Fig. 1C-D). However, resynthesis is probably minor for most vegetative proteins, since the housekeeping σ factor, σ^A^, shows low activity in forespores (Marquis et al. 2008).

To test if proteins are diluted by forespore growth, we suppressed forespore growth by two methods. First, we blocked growth with a *spoIIQ* mutation (Londoño-Vallejo et al. 1997; Sun et al. 2000; Broder and Pogliano 2006) and used fluorescence microscopy to assess the abundance of CitZ-GFP, Icd-GFP and GFP expressed from the *citZ* promoter (*P_citZ_*) in the forespore. The *spoIIQ* mutation had no impact on CitZ-GFP fluorescence in the forespore, with fluorescence decreasing to levels below detection (Fig 2A-B). However, *spoIIQ* forespores were smaller (Fig. S2) and contained similar levels of Icd-GFP and GFP fluorescence as the mother cell (Fig 2A-B). We also blocked forespore growth by inhibiting membrane synthesis with cerulenin (Lopez-Garrido et al. 2018; Landajuela et al. 2022), and measured the abundance of CitZ-GFP and Icd-GFP by timelapse fluorescence microscopy. Forespore fluorescence of CitZ-GFP decreased with nearly identical kinetics with or without cerulenin (Fig. 2C-D), whereas Icd-GFP fluorescence did not decrease in cerulenin-treated forespores. These results suggest that forespore growth mediates a widespread ∼4-fold reduction in FS:MC protein concentration, whereas a specific pathway mediates the dramatic depletion of CitZ from the forespore.

**Figure 2:**
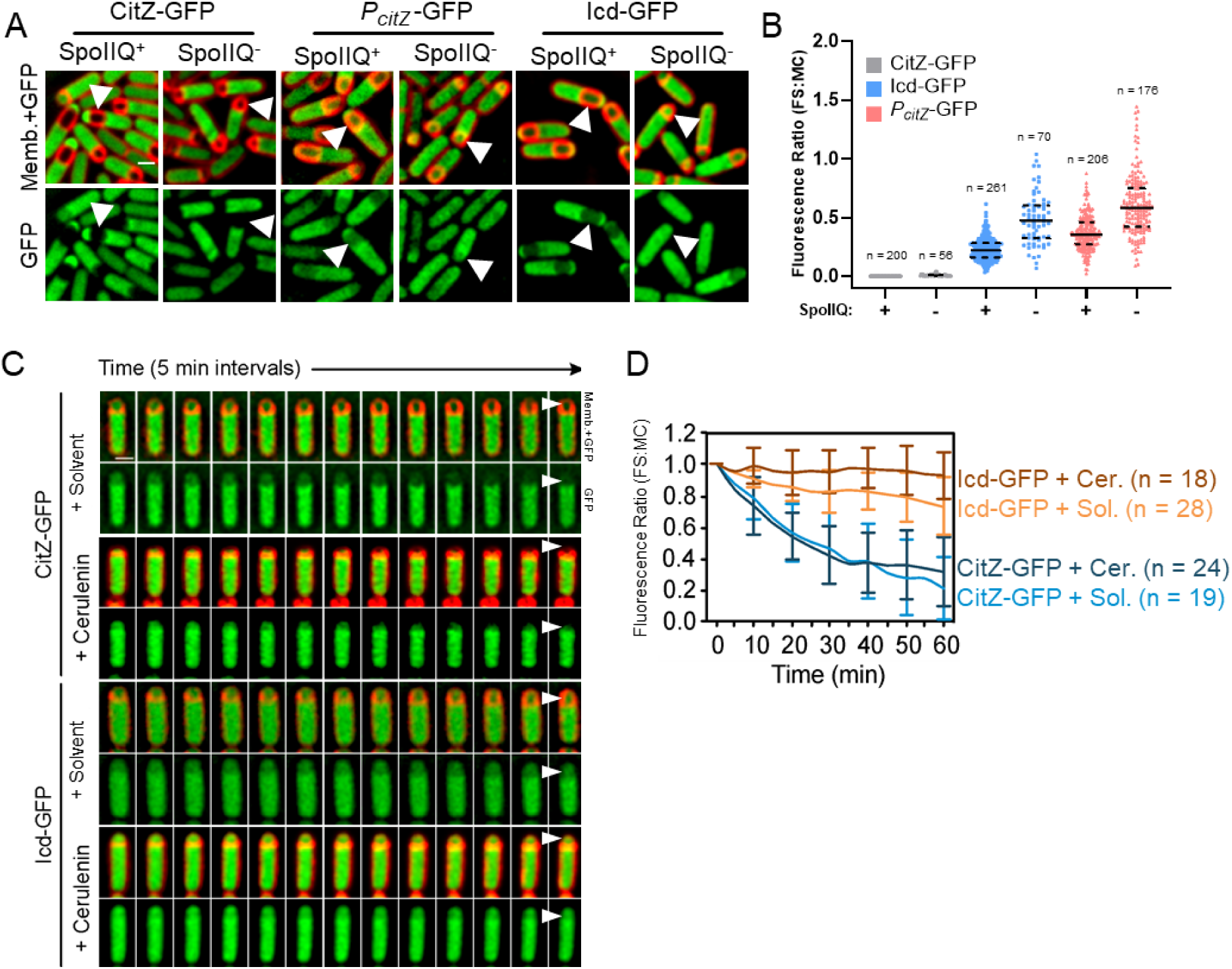
Forespore growth mediates protein dilution. (A) Visualization of CitZ-GFP and Icd-GFP (green), and from GFP produced from the *citZ* promoter (P*citZ*-GFP) in sporulating cells containing (SpoIIQ+) or lacking (SpoIIQ^−^) SpoIIQ. Arrowheads indicate forespores. Membranes are stained with FM 4-64 (red). (B) Quantification of the forespore-to-mother-cell (FS:MC) GFP ratio for the strains shown in (A). The “+” and “–” symbols on the *x*-axis indicate the presence and absence of SpoIIQ, respectively. Dashed lines indicate the middle 75% of the data and solid lines the median. (C) Time-lapse fluorescence microscopy of FM 4-64 stained sporulating cells producing CitZ-GFP (upper) or Icd-GFP (lower) in the presence of 30 µg/mL of cerulenin or an equal volume of solvent. Forespores are indicated by arrowheads on the last image. (D) Quantification of the FS:MC GFP ratio of sporulating cells from the timelapses shown in (B) over time, relative to the initial signal at t_0_. Values represent the mean ± SD. Scale bars, 1µm.

### The ClpCP protease responsible for CitZ depletion in the forespore

We hypothesized that the pathway leading to CitZ-GFP depletion involves proteolysis, since depletion is not explained by transcription (Fig. S3). We used a candidate gene approach to identify the protease, surveying a panel of null mutants in genes encoding proteases or protease accessory factors, including the AAA+ protease/protease components *clpC*, *clpE*, *clpQY, clpP, clpX, ftsH,* and *lon*, the carboxy terminal peptidases *ctpA* and *ctpB*, the intracellular serine protease *ispA*, and the germination protease *gpr*. Of the eleven null mutants tested, two failed to sporulate (*clpX* and *clpP*) so we could not test if they were involved in metabolic differentiation. Of the remaining nine, only the *clpC* mutation showed an appreciable stabilization of CitZ-GFP in the forespore (Fig. 3A-B). It is most likely that the absence ClpC stabilizes CitZ due to the inability to form a functional ClpCP protease, but we were unable to test if the *clpP* mutant stabilized CitZ due to its severe sporulation defect (Fig. 3A) (Msadek et al. 1998). To circumvent this limitation, we bypassed the sporulation initiation defect of the *clpP* mutant by inactivating phosphatases that reduce Spo0A phosphorylation (Nanamiya et al. 2000). CitZ-GFP was stabilized in the few ClpP^−^ forespores we were able to observe (Fig. S4A), indicating that ClpP, together with ClpC, is required for CitZ-GFP depletion.

**Figure 3.**
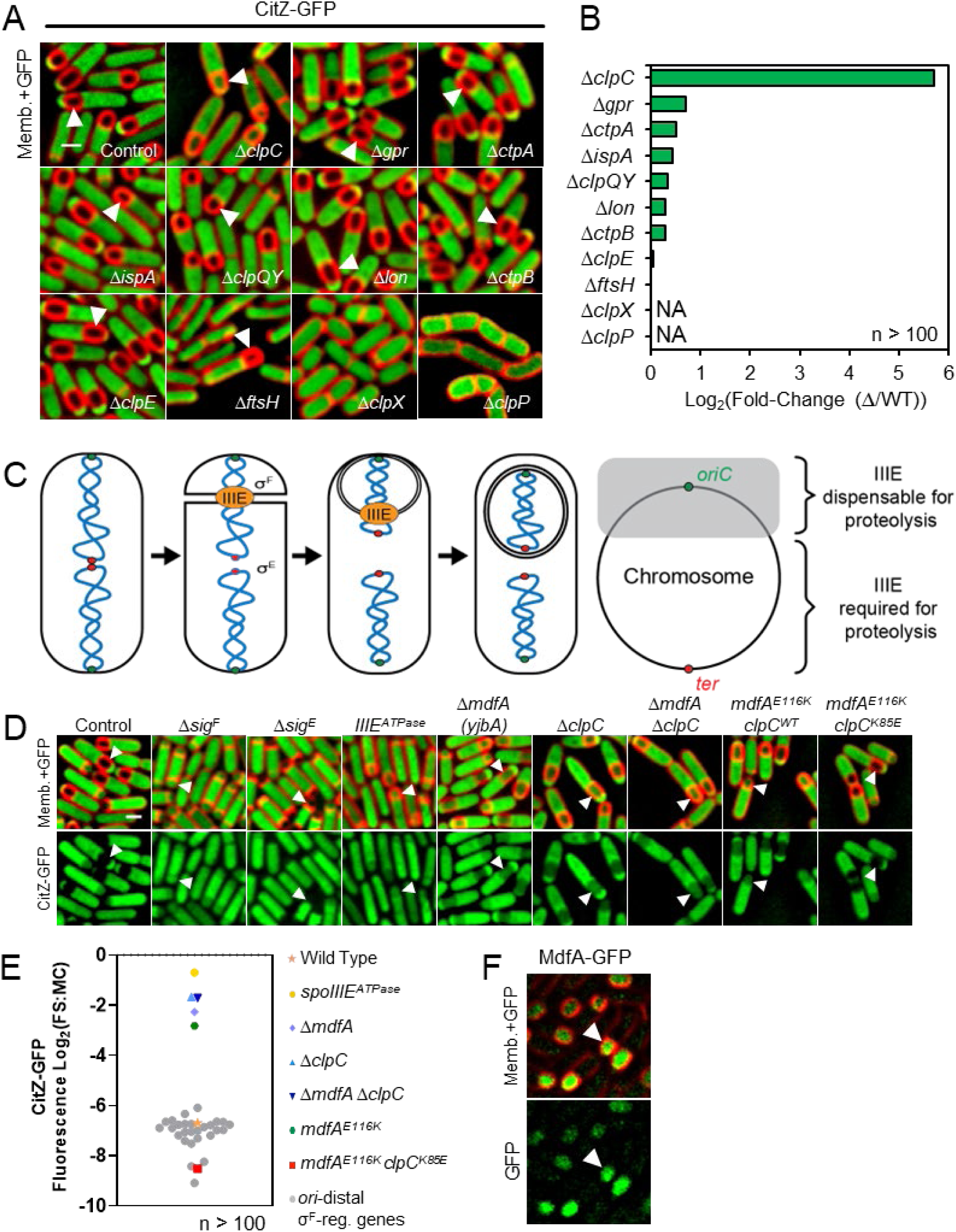
Identification of proteolytic pathway that mediates depletion of CitZ-GFP in the forespore. (A) Visualization of CitZ-GFP (green) in mutant sporulating cells lacking the indicated proteases. Membranes are stained with FM 4-64 (red). Scale bar, 1 µm. (B) Log_2_ of the fold change in the FS:MC GFP ratio for each strain relative to wild type. ClpX and ClpP mutants failed to sporulate, so it was not possible to calculate the CitZ-GFP fluorescence ratios (NA). More than one hundred cells were analyzed for each strain. (C) The newly synthesized polar septum traps the forespore chromosome (blue), with the origin of replication (*oriC*, green circle) in the forespore, and the terminus (*ter*, red circle) in the mother cell. The remaining forespore chromosome is translocated into the forespore by SpoIIIE (IIIE). The diagram on the right shows the chromosome highlighting the region initially trapped in the forespore (gray). IIIE is required to move genes in the unshaded region into the forespore. (D) Visualization of CitZ-GFP in sporulating cells lacking σ^F^ (Δ*sigF*), lacking σ^E^ (Δ*sigE*), containing a SpoIIIE allele unable to hydrolyze ATP (*IIIE^ATPase^*), lacking MdfA (Δ*mdfA*), lacking ClpC (Δ*clpC*), lacking both MdfA and ClpC (Δ*mdfA* Δ*clpC*), containing the MdfA E116K allele (*mdfA^E116K^ clpC^WT^*) or containing the MdfA E116K and the ClpC K85E alleles (*mdfA^E116K^ clpC^K85E^*). Membranes are stained with FM 4-64 (red). (E) Log_2_ of the FS:MC fluorescence ratio of each strain shown in (D) and of mutants lacking selected σ^F^-regulated genes initially trapped in the mother cell (*ori*-distal σ*^F^*-reg. genes). Every dot represents the Log_2_ of the average ratio of a different strain, color-coded according to the key at the right. More than one hundred cells were analyzed for each strain. (F) Localization of MdfA-GFP. Membranes are stained with FM 4-64 (red). Scale bars, 1 µm.

The ClpCP protease plays a critical role in cellular homeostasis and sporulation, and mutants in both *clpC* and *clpP* have pleiotropic effects (Msadek et al. 1998; Pan et al. 2001; Meeske et al. 2016). To minimize these effects, we used spatiotemporally regulated proteolysis (STRP) to specifically deplete ClpC-ssrA* after σ^F^ activation (Fig. S4B) (Riley et al. 2018). Depletion of ClpC-ssrA* in the forespore resulted in the marked stabilization of CitZ-GFP (Fig. S4C-G), suggesting that the impact of the *clpC* mutation on CitZ-GFP levels is due to the direct action of ClpCP.

### MdfA is required for forespore-specific degradation of CitZ

We previously noted that CitZ-GFP degradation required the early forespore transcription factor σ^F^ (Riley et al. 2021a). The *clpC* gene is part of the σ^F^ regulon (Wang et al. 2006) and ClpC becomes enriched in the forespore, which could confer cell-specificity to degradation (Kain et al. 2008). However, when *clpC* and *clpP* were expressed from inducible promoters, CitZ-GFP remained stable in the mother cell (Fig. S4F-G), suggesting that another mechanism, likely involving a protein whose expression depends on σ^F^, confers forespore-specific CitZ degradation. To identify this protein, we took advantage of the transient genetic asymmetry between the mother cell and forespore after polar septation (Wu and Errington 1998). Specifically, the newly-synthesized sporulation septum traps the forespore chromosome, with the origin-proximal 1/3 of the chromosome initially located in the forespore and the remaining 2/3 translocated into the forespore by the SpoIIIE ATPase (Fig. 3C) (Wu and Errington 1994; Ling Juan et al. 1995; Bath et al. 2000). Cells expressing mutant SpoIIIE proteins lacking ATPase activity activate σ^F^ and compartmentalize forespore gene expression but fail to translocate the origin-distal region of the chromosome into the forespore (Sharp and Pogliano 1999). Therefore, if the gene encoding the unknown metabolic differentiation factor is located at an origin-proximal locus, then forespore-specific CitZ degradation would not require SpoIIIE activity. In contrast, if the gene is on the origin-distal portion of the chromosome, then forespore-specific CitZ degradation would depend on SpoIIIE activity. To test this hypothesis, we introduced a SpoIIIE ATPase mutation into strains expressing CitZ-GFP and found that CitZ-GFP was stabilized in the forespore (Fig. 3D-E), suggesting the gene encoding the unknown differentiation factor lies in the origin-distal part of the chromosome.

We next identified 28 genes that were regulated by σ^F^, located in the origin-distal 2/3 of the chromosome and encoded a protein of unknown function. We obtained null mutants of 28 candidate genes meeting these criteria from the *Bacillus* Genetic Stock Center (Koo et al. 2017) and used fluorescence microscopy to screen for mutants that stabilized CitZ-GFP in the forespore to a similar extent as the *clpC* or *sigF* null strains (Fig. 3D). One null mutation *(yjbA)* stabilized CitZ-GFP (Fig. 3D-3E; Fig. S5). Based on results presented later here and in the accompanying paper reporting the independent identification of the gene as encoding a forespore-specific ClpC adaptor protein (Massoni et al. 2024), we renamed the *yjbA*-encoded protein MdfA, for metabolic differentiation factor A, and the gene *mdfA*. We found that the null mutant was able to complete spore assembly (Fig. 4C), and that MdfA-GFP localized to the forespore cytoplasm (Fig. 3F).

**Figure 4.**
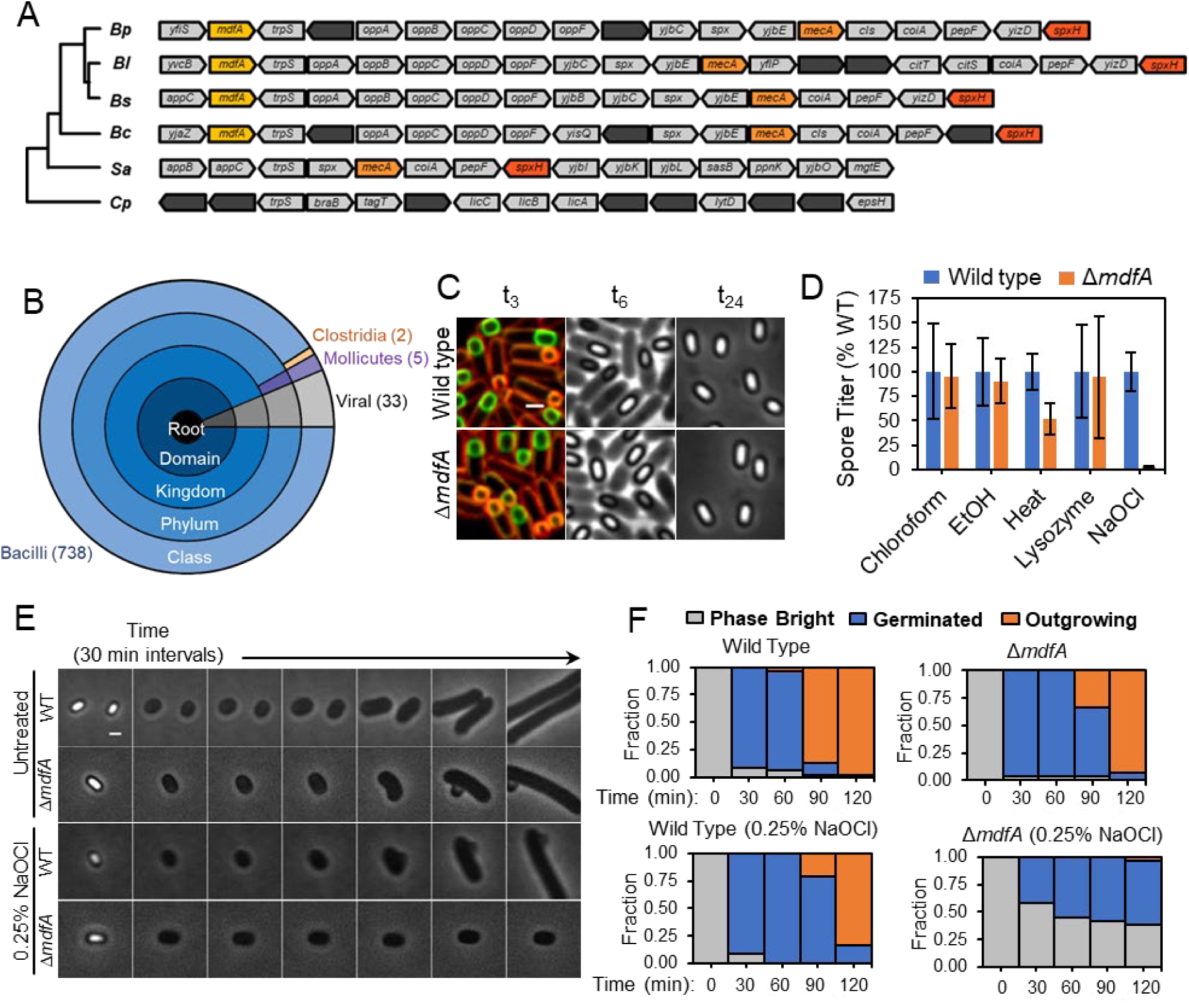
MdfA protects spores from oxidative damage. (A) Phylogenetic tree of *B. pumilus* (*Bp*), *B. licheniformis* (*Bl*), *B. subtilis* (*Bs*), *B. cereus* (*Bc*) *Staphylococcus aureus* (*Sa*) and *Clostridium perfringens* (*Cp*). The *mdfA* region is shown for every species, with each arrow representing a gene. *mdfA*, yellow. Other ClpC adaptor proteins include *mecA*, orange; *spxH*, red. (B) Sunburst diagram from PFAM representing the distribution of MdfA homologs across different taxa, with the majority being found in the *Bacilli* (blue), with a few in viruses (gray), *Mollicute*s (purple), *Clostridia* (orange). (C) Sporulating cultures of the wild type and the Δ*mdfA* mutant at t_3,_ t_6_ and t_24_ after resuspension. Membranes were stained with FM 4-64 (red) and MitotrackerGreen (green) for t_3_ samples, to discriminate between partially engulfed forespores (orange) and fully engulfed spores (green). Phase-contrast images are shown at t_6_ and t_24_. (D) Relative spore titers of the wild-type (blue) and Δ*mdfA* (orange) strains after nutrient exhaustion and treatment with chloroform, ethanol (EtOH), heat, lysozyme, or NaOCl. Wild-type titers were normalized to 100%. Values represent the mean±SD of three independent experiments. (E) Phase-contrast timelapse microscopy of reviving spores from the wild-type and Δ*mdfA* strains treated with 0.25% NaOCl for 18 min or untreated. Images were acquired every 30 min for 3.5 h. (F) Quantification of the fraction of phase bright (grey), germinated (blue), and outgrowing (orange) spores during timelapse experiments. More than one hundred spores were analyzed for each strain and treatment. Scale bars, 1 µm.

### The interaction between MdfA and the ClpCP protease is required for CitZ depletion

The accompanying manuscript demonstrates that MdfA and ClpC interact, and that MdfA promotes ClpC assembly and ATPase activity, suggesting that MdfA is a ClpCP adapter protein (Massoni et al. 2024). To determine if this interaction is required for forespore-specific degradation of CitZ, we made use of the co-crystal structure from Massoni, Evans *et al*., which predicts that MdfA and ClpC interact in part via a salt bridge between E116 in MdfA and K85 in ClpC. A single amino acid change that abolished this interaction (MdfA^E116K^) stabilized CitZ in the forespore (Fig. 3D-E). Introducing a second mutation that restored the interaction (ClpC^K85E^) restored forespore-specific degradation of CitZ in vivo (Fig. 3D-E). In addition, the *clpP* mutation stabilized CitZ (Fig. S4A). Thus, our data and that of Massoni, Evans *et al*. indicate that MdfA is a forespore-specific ClpCP adaptor that promotes degradation of CitZ in the forespore.

### Metabolic differentiation is required for spore resistance to the oxidant hypochlorite

Genes encoding MdfA homologs are broadly distributed among the aerobic spore forming *Bacilli* but absent from the genomes of non-sporulating bacteria and the anaerobic spore forming *Clostridia* (Fig. 4A-B), so we hypothesized that MdfA activity might be important in overcoming environmental challenges faced uniquely by *Bacillus* spores. The *ΔmdfA* strain produced phase bright spores that appear similar to wild type (Fig. 4C), suggesting that the mutant does not prevent spore assembly. These spores show normal resistance to chloroform, ethanol and lysozyme, a marked 20-fold decrease in resistance to hypochlorite treatment and a 2-fold decrease in heat resistance (Fig. 4D); the latter could be due to heat causing a secondary oxidative stress (Mols and Abee 2011). The few colonies formed by hypochlorite-treated *DmdfA* spores were smaller those formed by wild-type, so we performed time lapse microscopy to investigate germination and outgrowth at the single cell level. Consistent with population level observations, untreated *DmdfA* spores transitioned from phase bright to phase dark and initiated outgrowth with the same kinetics as wild type (Fig. 4E-F), whereas hypochlorite treated *DmdfA* spores showed severely delayed germination and outgrowth. This suggests that MdfA is required to produce a spore that is able to germinate and resume growth after oxidative stress induced by hypochlorite.

### Identification of proteins depleted from the spore in an MdfA-dependent manner

To assess the impact of the MdfA/ClpCP pathway on the spore proteome, we used mass spectrometry to identify proteins that are depleted from the forespore in an MdfA-dependent manner. We isolated spores from MdfA^+^ and MdfA^−^ strains, extracted their proteins and determined the relative amounts of individual proteins in the two backgrounds using quantitative proteomics (Fig. 5A). To facilitate extraction of spore proteins, we germinated the spores briefly in the presence of chloramphenicol to block protein synthesis and used strains lacking the germination proteases Gpr and TepA (Sussman and Setlow 1991; Traag et al. 2013).

**Figure 5.**
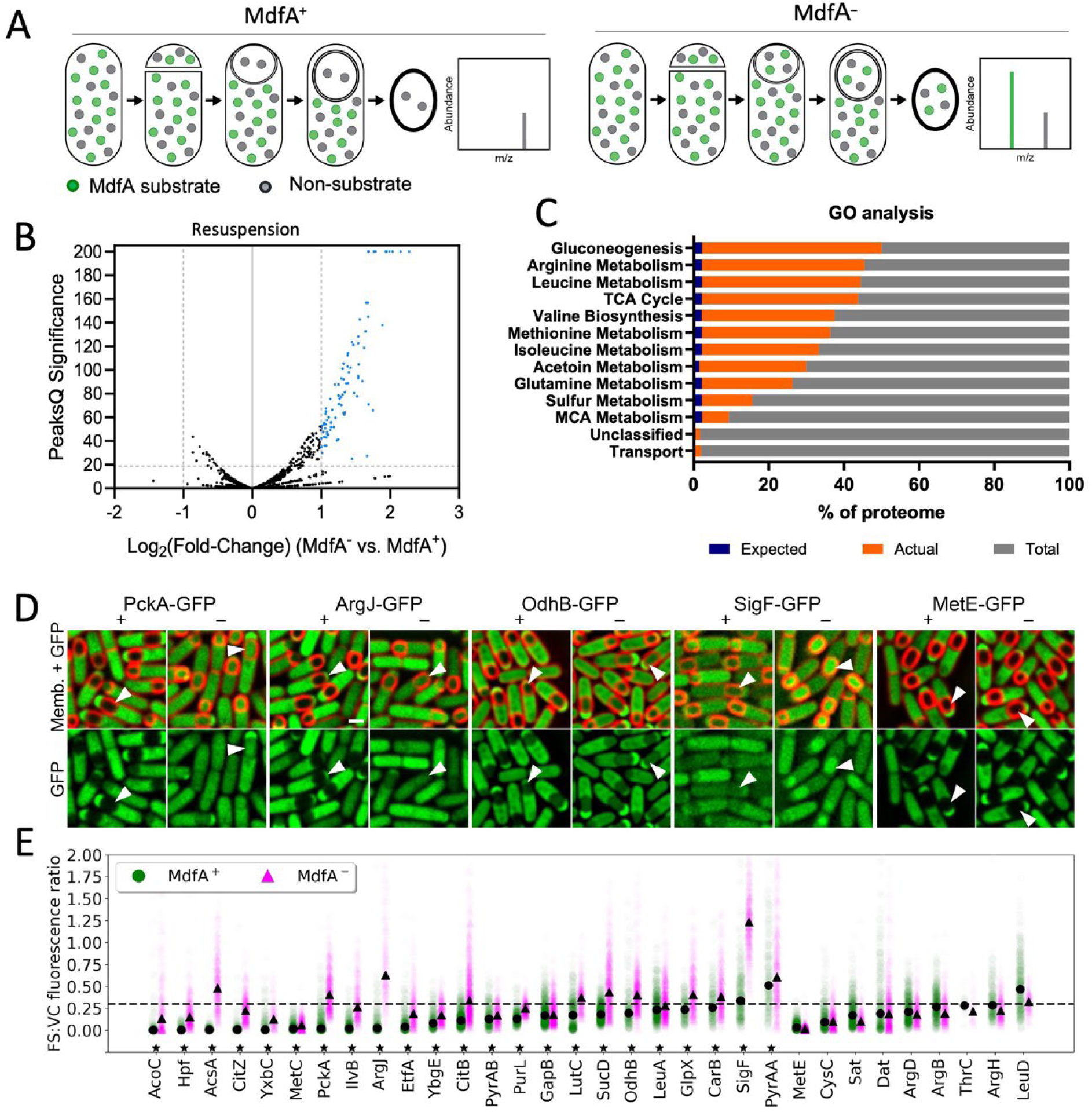
Identification of proteins depleted in a MdfA-dependent manner. (A) Strategy to identify proteins depleted from the spore in a MdfA-dependent manner. Non MdfA substrates (gray circles) are present in similar amounts in both MdfA^+^ and MdfA*^−^* spores. MdfA substrates (green) are more abundant in MdfA^−^ than in MdfA^+^ spores. (B) Volcano plot of mass-spectrometry results comparing protein abundance in MdfA^−^ vs. MdfA^+^ spores when sporulation was induced by resuspension (see methods). Proteins with >1-Log_2_ fold change and >20 significance were considered potential substrates (blue dots). 117 hits were observed (Data Table 1, with raw data available via ProteomeXchange (Vizcaino et al. 2013), identifier PXD051727). (C) Gene Ontology analysis of the MdfA substrates identified through proteomics based on PANTHER results (Data Table 1). Blue indicates the representation of each pathway in the *B. subtilis* proteome and the expected representation of the pathway, orange indicates the actual representation of proteins in each pathway in the subset of proteins that were enriched in the absence of MdfA. (D) Examples of validation using GFP fusions. (full set in Fig. S7). GFP fusion proteins (green) were visualized in MdfA^+^ (+) and MdfA^−^ (–) sporulating cells stained with FM 4-64 (red). Arrowheads indicate forespores. Scale bar, 1 µm. (E) Forespore (FS) GFP fluorescence relative to the median fluorescence of vegetative cells (VC) present in the same sporulating cultures of strains carrying GFP fusions to the indicated proteins in MdfA^+^ (green circles) and MdfA^−^ (magenta triangles) strains. The black dots and triangles indicate the median of each series; every green circle and magenta triangle represent the relative fluorescence of an individual forespore. >150 forespores were analyzed for each protein (except for ThrC, where 23 and 44 forespores were analyzed in MdfA^+^ and MdfA^−^ backgrounds, respectively). The horizontal dashed line marks relative forespore fluorescence value of 0.3, expected if vegetative proteins were only diluted by forespore growth. The asterisks by the protein names indicates those that were significantly enriched in MdfA^−^ forespores (P<0.01). These are shown first, sorted from lowest to highest MdfA^+^ FS:VS fluorescence ratio. Non significantly enriched proteins are shown last, sorted in the same way.

We identified 117 proteins that were enriched in MdfA^−^ spores compared to MdfA^+^ spores (Fig. 5B, Data Table S1), with 42 found in both sporulation conditions, exhaustion and resuspension. Metabolic proteins represented 90% of the proteins present at higher levels in the MdfA^−^ spores, with proteins involved in gluconeogenesis, the TCA cycle (including CitZ), monocarboxylic acid metabolism and synthesis of the amino acids arginine, leucine, valine, isoleucine, and glutamine significantly overrepresented among the hits relative to the proteome (Fig. 5C). We also observed elevated levels of the transcription factors σ^F^, σ^E^, σ^B^, RsbW (which inhibits σ^B^ activity (Benson and Haldenwang 1993)), ribosome hibernation promoting factor Hpf (Akanuma et al. 2016), and the cell division proteins FtsZ and DivIVA (Data Table 1). There is strong agreement between these proteins and those found to be less abundant in spores than growing cells in proteomic experiments (Swarge et al. 2020; Huang et al. 2024), with almost 90% of the proteins here identified as being depleted in an MdfA-dependent manner also being reported to be deficient in mature spores by Swarge *et al*. (Fig. S6).

To confirm these results, we created C-terminal GFP fusions to dozens of the proteomic hits and determined if the fusion proteins were stabilized in MdfA^−^ forespores. To facilitate analysis, we focused on 32 cytoplasmic fusion proteins (Fig. 5E; Fig. S7). We quantified forespore and mother cell fluorescence, and compared these values to vegetative cell fluorescence in the same samples. As expected, the effect of MdfA was much more dramatic in forespores than mother cells (Fig. S8). We found that 23 fusion proteins showed a statistically significant increase in GFP fluorescence in MdfA^−^ forespores compared to wild type (Fig. 5E). Some of these proteins were dramatically depleted in wild type forespores and strongly stabilized in MdfA^−^ forespores (Fig. 5E), suggesting they may be direct targets of the MdfA/ClpCP. These include acetyl-CoA synthetase (AcsA), gluconeogenic protein phosphoenolpyruvate carboxykinase (PckA), arginine biosynthetic protein N-acetylglutamate synthase (ArgJ), and ribosome hibernation factor (Hpf), among others (Fig. 5D-E; Fig. S7). Other proteins showed variable levels of depletion in wild type forespores and modest but statistically significant enrichment in MdfA^−^ forespores (Fig. 5D-E). These proteins may be low-affinity substrates of MdfA/ClpCP, or they could be indirectly stabilized in the absence of MdfA. Interestingly, the early forespore sigma factor σ^F^ (SigF-GFP) showed no evidence of depletion in wild type forespores, but was present at ∼4-fold higher levels in MdfA^−^ forespores (Fig. 5D, 5F; Fig. S8A-B), suggesting that the increased level of σ^F^ in MdfA^−^ forespores is in part due to increased σ^F^ synthesis. The 9 proteins that showed no statistically significant difference between the fluorescence of MdfA^+^ and MdfA^−^ forespores (Fig. 5E) could be false-positive proteomics hits or the C-terminal GFP fusion could interfere with MdfA-mediated degradation. Interestingly, we also identified a few proteins that appeared to be depleted from the forespore in an MdfA-independent manner, such as MetE-GFP (Fig. 5D-E), suggesting the existence of a separate forespore specific degradation pathway.

## Discussion

We previously demonstrated that *B. subtilis* sporulation entails the marked metabolic differentiation of the two cells required to produce a spore, with the developing spore losing the capability to synthesize metabolic precursors and thereby becoming dependent on mother cell metabolites to support the protein synthesis needed to complete development (Riley et al. 2021a). We here identify a proteolytic pathway, comprised of the forespore-specific protein MdfA (YjbA), the unfoldase ClpC and the ClpP protease, that actively depletes proteins from the forespore and contributes to metabolic differentiation (Fig. 6A-B). MdfA was independently identified in the companion manuscript as a ClpC adapter protein that stimulates ClpC ATPase activity (Massoni et al. 2024). We characterize the impact of the MdfA/ClpC pathway on the spore proteome, and find that it mediates the depletion of a highly specific subset of target proteins, most of which are involved in metabolic processes such as gluconeogenesis, the TCA cycle, and amino acid synthesis, with some involved in cell division and gene expression. Thus, metabolic differentiation severely limits the metabolic pathways that can function in the developing spore, necessitating forespore nurturing by the mother cell and the transport of nutrients via the Q-A channel and potentially other unidentified channels (Camp and Losick 2009; Riley et al. 2021a; Tibocha-Bonilla, Lyda et al., in revision).

**Figure 6.**
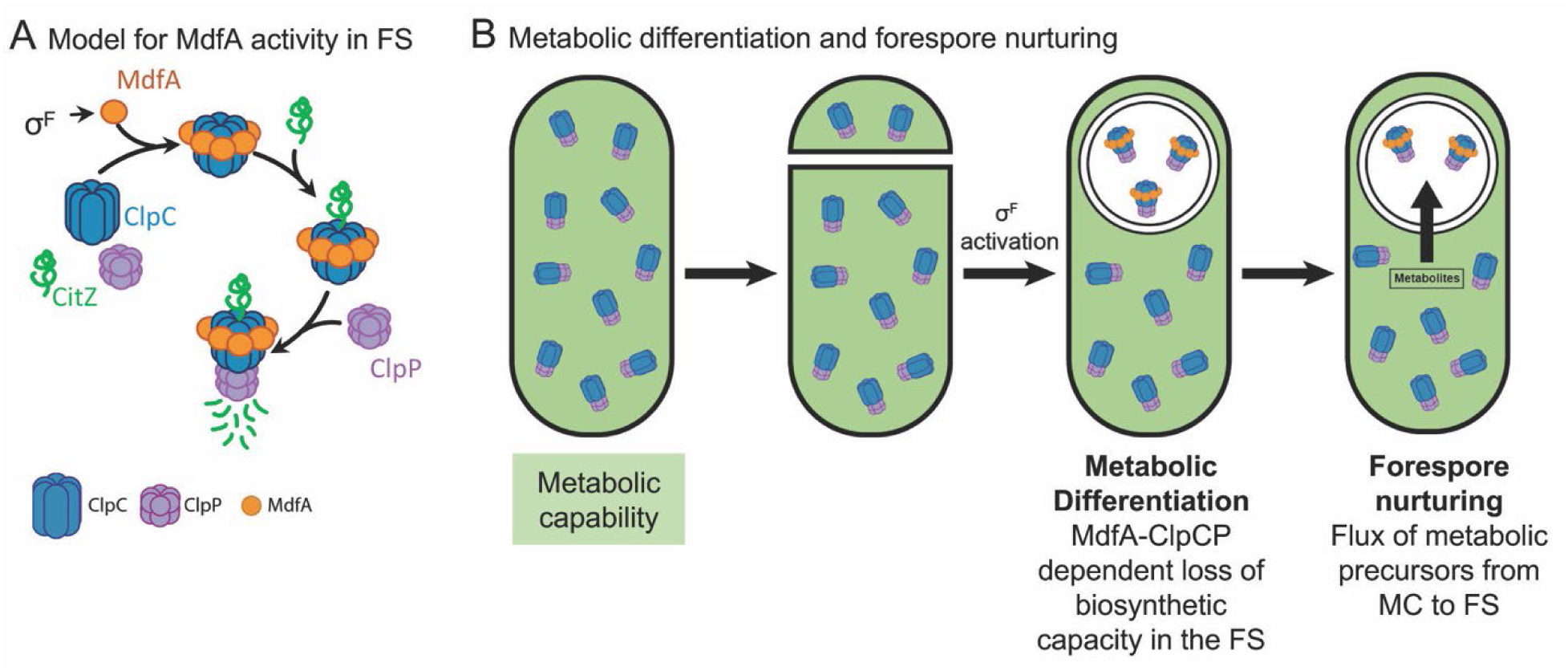
Model of MdfA-dependent metabolic differentiation. (A) MdfA (orange) is produced in the forespore under σ^F^ control, interacts with ClpC (blue), and delivers target proteins (green) to ClpP (purple) for proteolysis. (B) In vegetative cells, ClpC and ClpP participate in protein quality control. During sporulation, MdfA is produced in the forespore, delivering target proteins to ClpCP for degradation, reducing the metabolic capacity of the forespore (green shading), which becomes dependent on the mother cell for metabolic precursors for biosynthesis.

Proteolysis is widely used in *B. subtilis* to mediate the transition between growth and different developmental states, including competence (Turgay et al. 1998), biofilm formation (Mukherjee et al. 2015), and sporulation (Pan et al. 2001; Bradshaw and Losick 2015). Each of these pathways typically deploys specific adaptor proteins to cause the degradation of cell-fate determinants or transcriptional regulators, thereby achieving differentiation indirectly through the modulation of transcription (Kirstein et al. 2009). The MdfA/ClpCP proteolytic pathway may play a similar role during sporulation, as it governs the levels of several transcription factors, such as σ^F^, σ^E^ and σ^B^, in the forespore (Data Table 1). This may reinforce developmental gene expression by eliminating σ factors that compete with the forespore-specific σ factors for binding to the RNA polymerase (Haldenwang 1995; Collins et al. 2023). It may also contribute to the progression from early σ^F^-dependent to late σ^G^-dependent forespore-specific gene expression by limiting σ^F^-activity, as noted in Massoni, Evans et al. (2024). Interestingly, the ClpCP protease was previously observed to reinforce the activation of σ^F^ by depleting the anti-σ factor SpoIIAB from the forespore (Pan et al. 2001). SpoIIAB was not identified in proteomic experiments, suggesting that it may be degraded in an MdfA-independent manner, or that there is a redundant proteolytic pathway.

The increased levels of some proteins in the absence of MdfA are not completely explained by proteolysis. For example, the early forespore σ factor σ^F^ is present in wild-type forespores at levels predicted by dilution, but levels increase in the absence of MdfA, suggesting ongoing synthesis in the absence of MdfA (Fig. 5E). It is possible that MdfA reduces σ^F^ synthesis in a proteolysis-independent manner by interfering with *sigF* transcription, as proposed for the inhibition of sporulation initiation and biofilm formation by the ClpC adaptor protein MecA (Prepiak et al. 2011; Tanner et al. 2018). However, we favor the hypothesis that σ^F^ is an MdfA-ClpCP target that, unlike most metabolic enzymes, continues to be synthesized in the forespore from the σ^F^-dependent *dacF* promoter, which has been shown to cause read-through transcription of *sigF* in vivo (Schuch and Piggot 1994; Nicolas et al. 2012). In this model, the low-level persistence of σ^F^ in wild-type forespores would reflect the balance between synthesis and degradation, whereas the higher-than-expected levels of σ^F^ in MdfA-forespores would be caused by decreased σ^F^ proteolysis, resulting in increased σ^F^ levels and increased transcription of *sigF* from the *dacA* promoter. Regardless of the mechanism, we hypothesize that preventing continued synthesis of σ^F^ in the forespore facilitates the replacement of σ^F^ by σ^G^ and progression through the developmental gene expression cascade.

Our results suggest that the transition of the forespore to dormancy starts shortly after polar septation, with the synthesis of MdfA and the activation of MdfA/ClpCP proteolysis. Spores that fail to undergo this transition due to the absence of MdfA are highly sensitive to hypochlorite-induced oxidative stress. Previous work has demonstrated that germination and outgrowth place a tremendous oxidative burden on the reviving spore (Ibarra et al. 2008), likely due to the production of reactive oxygen species during metabolic reactivation. The electron transport chain is a major source of reactive oxygen species (Zhao et al. 2019), and we hypothesize that metabolic differentiation protects germinating spores against oxidative damage by reducing the levels of dehydrogenases and TCA cycle enzymes to decrease flux through the electron transport chain. This is consistent with prior data demonstrating that, although spores accumulate large stores of the glycolytic intermediate 3-phosphoglycerate that could readily feed into the TCA cycle and fuel electron transport (Nelson and Kornberg 1970), germinating spores do not resume oxidative metabolism until late in outgrowth after stress responses have been activated (Sinai et al. 2015; Swarge et al. 2020). Metabolic differentiation might therefore reduce the oxidative burden of metabolism until the newly germinated cell is fully equipped to handle stress.

The mechanism by which MdfA/ClpC targets proteins for degradation remains unclear. Our proteomics and quantitative fluorescence microscopy data indicate that dozens of proteins are stabilized in MdfA^−^ forespores. It is possible that many of these proteins contain structural features that are directly recognized by the MdfA/ClpC pathway. However, it is also possible that proteins stabilized in MdfA^−^ forespores are indirectly targeted. In this second scenario, the MdfA/ClpC pathway might recognize only a few substrates located at key metabolic junctions (Tibocha-Bonilla et al. 2022; Tibocha-Bonilla, Lyda *et al*. in revision), so that their elimination would block flux through the pathways in which they participate, rendering downstream enzymes inactive. Inactive enzymes would then be degraded by ClpCP in an MdfA-independent manner (Kruger et al. 2000; Gerth et al. 2008; Gerth et al. 2017). This model is consistent with the observations that ClpCP degrades enzymes that are no longer needed (Gerth et al. 2017) and that MdfA activates the ATPase activity of ClpC (Massoni et al. 2024), suggesting that it enhances general proteolysis by ClpCP in the forespore. Interestingly, we have observed that CitZ proteins with mutations in the active site are unstable *in vivo* (not shown), suggesting that they are targeted for proteolysis. Clearly, further work is needed to decipher the targeting mechanism and to identify the direct and indirect targets of the MdfA/ClpCP pathway.

Our results support those presented in the companion manuscript by Masoni, Evans *et al*., which show that MdfA is the founding member of a new sporulation-specific family of ClpCP adaptor proteins. Together with the mother-cell ClpXP adaptor CmpA, which mediates SpoIVA degradation when the spore envelope is misassembled (Tan et al. 2015), the production of MdfA in the forespore illustrates how Clp proteolysis is exploited during sporulation through the cell-specific expression of adaptor proteins to remodel the proteome and mediate differentiation. Collectively, the results of the two manuscripts demonstrate that MdfA-induced proteolysis is a newly recognized aspect of sporulation that mediates metabolic differentiation and contributes to cell-specific gene expression during spore formation, making sporulation one of the few examples where metabolic reprogramming of genetically identical sister cells is achieved by proteolysis (Wood and Haselkorn 1980; Mutomba and Wang 1998). Further studies are required to document the full impact of MdfA/ClpCP proteolysis on the formation, resistance properties, and revival of bacterial endospores.

## Materials and Methods

### Strain Construction

Strains are derivatives of *B. subtilis* PY79 (Table S1). Strains were constructed by transformation using plasmids and oligonucleotides in Table S2 and S3 as described in supplementary materials. Antibiotic concentrations for *B. subtilis* were 10 µg/mL kanamycin, 100 µg/mL spectinomycin, 5 µg/mL chloramphenicol, 10 µg/mL tetracycline, 1 µg/mL erythromycin and 25 µg/mL lincomycin.

### Culture Conditions

For fluorescence microscopy, sporulation was induced by resuspension (Sterlini and Mandelstam 1969) modified as described (Riley et al. 2021a) at 37°C for batch culture and 30°C for timelapse microscopy. For spore titers and spore purification, sporulation was induced by nutrient exhaustion in Difco Sporulation Medium (DSM) at 37°C (Schaeffer et al. 1965). Expression from P*_xylA_* and P*_spank_* was induced using 1% of xylose and 1 mM of IPTG, respectively.

### Microscopy and image analysis

Cells were visualized on an Applied Precision DV Elite or Ultra optical sectioning microscope equipped with a Photometrics CoolSNAP-HQ2 or a PCO Edge sCMOS camera. Images were deconvolved using SoftWoRx and quantitative image analysis was performed as described in (Riley et al. 2021a). Median focal planes are shown. Images in Fig. S4A were taken with a Leica THUNDER Imager Live Cell with a Leica DFC9000 GTC sCMOS camera, and computationally cleared using LAS X.

### Batch Culture Microscopy

Membranes were stained with 0.5 μg/mL FM 4-64 one hour after induction of sporulation by resuspension (*t_1_*). At *t_3_* (three hours after sporulation induction), 12 μl of culture was transferred to 1.2% agarose pads prepared in A+B medium and supplemented with an additional 0.5 μg mL^−1^ FM 4–64 (Life Technologies). Excitation/emission filters were TRITC/CY5 for membranes and FITC/FITC for GFP.

### Fluorescence Timelapse Microscopy

Fluorescence timelapse microscopy was performed as previously described (Riley et al. 2021a). Briefly, at *t_1_* cultures were stained with 0.5 μg/mL FM 4-64, and at t3 samples were collected and applied to an agar pad. Images were collected every 5 min for at least 2h. Sporangia were aligned vertically (with forespore on top) using Photoshop.

### Testing spore resistance

To test spore resistance, sporulation was induced by nutrient exhaustion in 2 mL of DSM for 24 hours at 37°C, and the cultures subject to various treatments, then serially diluted in 1x T-Base, and plated on LB. Heat resistance was tested by heating cultures at 80°C for 20 min, lysozyme resistance by incubating the cultures with 0.25 mg/mL lysozyme for 10 min at 37°C, chloroform and ethanol resistance by incubating cultures with 1:10 volumes of chloroform or 90% ethanol for 10 min at room temperature, and NaOCl resistance by incubating cultures with 0.25 % NaOCl for 18 min at room temperature. Spore titers were calculated based on the number of colony forming units (CFU) after overnight incubation at 30°C.

### Phase Contrast Timelapse Microscopy of Germination

Spores were purified from DSM cultures after 72 hours of incubation at 37°C, using a phosphate-polyethylene glycol aqueous biphasic gradient as previously described (Harrold et al. 2011; Riley et al. 2021a), and diluted to 0.3 O.D._600_ in sterile water. Phase contrast timelapse microscopy was performed on 1.2% agarose pads in LB plus 10 mM L-alanine (Riley et al. 2018). Quantification used spores that were phase-bright in the first timepoint. To test the impact of NaOCl, purified spores were treated with 0.25 % (v/v) NaOCl for 18 min, pelleted by gentle centrifugation, and washed 3 times with phosphate-buffered saline prior to imaging.

### Preparation of spores for mass spectrometry

Proteomics experiments were performed in strains lacking the two major germination proteases, *gpr* and *tepA* to limit protein turnover during germination. Sporulation of MdfA^+^ and MdfA^−^ strains was induced by resuspension in A+B medium or by nutrient exhaustion in DSM at 37°C. After 72 hours, cultures were pelleted and washed once with 4°C sterile water and incubated overnight at 4°C in sterile water to lyse vegetative cells. Spores were purified as described (Riley et al. 2021a). To facilitate protein extraction, germination was induced (Harwood et al. 1990). Briefly, spores were heat activated at 70°C for 20 min in water, resuspended in 10 mM Tris-HCl, pH 8.4 and diluted an O.D.580 of 0.3. Samples were incubated at 37°C for 20 min, and germination was induced by the addition of L-alanine 10 mM in the presence of 30 µg/mL chloramphenicol to block protein synthesis. Approximately 95% of the spores lost heat resistance by 15 min, at which time samples were harvested by centrifugation. The resulting spore pellets were stored at -80°C.

### Proteomic sample preparation

MS samples were prepared from purified spores and analyzed by the University of California, San Diego Biomolecular and Proteomics MS Facility (http://bpmsf.ucsd.edu/), as described in the supplemental materials.

## Acknowledgments

This work was supported by National Institutes of Health Grants (R01 GM57045 and R01 AI-113295 to KP; DP2 GM105439 to AHC; T32 GM007240-37 for EPR and JAL), the European Research Council (Starting Grant 853383 to JLG), the Biotechnology and Biological Sciences Research Council (BB/N006267/1, BB/R006091/1, BB/S006877/1, and BB/X001415/1 to RLI) and the University of California-HBCU initiative (for JAL). We thank Richard Losick for his helpful suggestions.

## Author Contributions

EPR, JLG, KP designed the experiments, EPR, JAL, IRK, EE, EA, SR performed the experiments, JS, ORM developed software, EPR, JLG, JAL, KP drafted the manuscript, all authors contributed to the final editing, and KP, AC, RIL and JLG aquired funding.

