## Supplementary information for "Developmentally-regulated proteolysis by MdfA and ClpCP mediates metabolic differentiation during *Bacillus subtilis* sporulation"

**Supplementary information for Riley, Lyda, et al**

**Contents**

- Figures S1 to S8

- Tables S1 to S3

- Details of plasmid construction

-Details of proteomics sample preparation

Supplementary Data Table S1 (separate Excel spreadsheet)

**Supplementary Data**

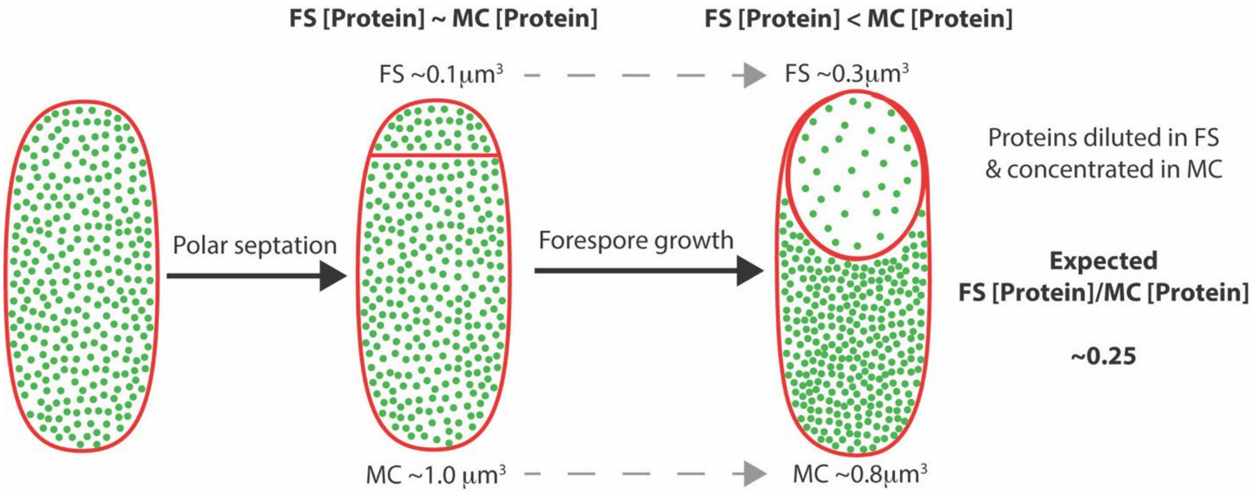


**Figure S1. Forespore growth causes proteins to be diluted in the forespore and concentrated in the mother cell.** Immediately after polar septation, the volume of the forespore is approximately one tenth of that of the mother cell (0.1 $\mu m^3$ vs. 1 $\mu m^3$), and cytoplasmic proteins (green dots) are equally concentrated in the two cells. As sporulation progresses, the forespore volume triples in detriment of the mother cell volume (Lopez-Garrido et al. 2018). This causes pre-existing cytoplasmic vegetative proteins to be diluted ~3 times in the forespore, and concentrated ~1.25 times in the mother cell. As a consequence, the concentration of cytoplasmic proteins becomes around 4 times higher in the mother cell than in the forespore. Differential synthesis or degradation of specific proteins in the two cells could lead to deviations of the expected FS:MC ratio.

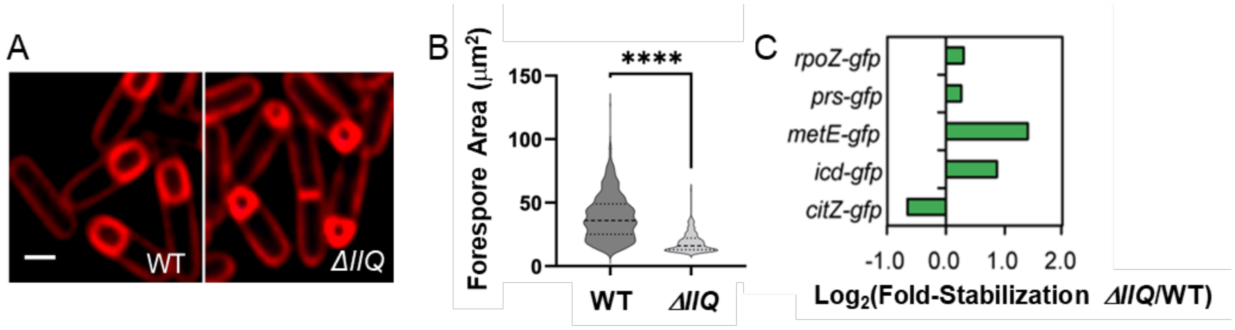


**Figure S2. Strains lacking SpoIIQ are impaired in forespore growth.** (A) Fluorescent microscopy of wild-type (WT) and Δ*spoIIQ* (ΔIIQ) sporulating cells. Membranes were stained with FM 4-64 (red). Scale bar, 1 μm. (B) Area of wild-type (WT) and Δ*spoIIQ* (*ΔIIQ*) forespores. Dotted lines in the violin plots delimit the middle 75% of the data, and the dashed lines represent the median. More than one hundred forespores were measured for every strain. The difference between the two strains is statistically significant ($P < 0.0001$) (C) Bar graph showing stabilization of GFP fusions to different metabolic enzymes in the forespores of Δ*spoIIQ* strains. The x-axis compares the forespore-to-mother-cell fluorescence ratio of each fusion in a Δ*spoIIQ* genetic b ackground to the respective forespore to mother cell fluorescence ratio in the wild-type background and represents the $\log_2(FS^{\Delta spoIIQ}:MC^{\Delta spoIIQ}/FS^{WT}:MC^{WT})$. More than one hundred cells were analyzed for each strain.

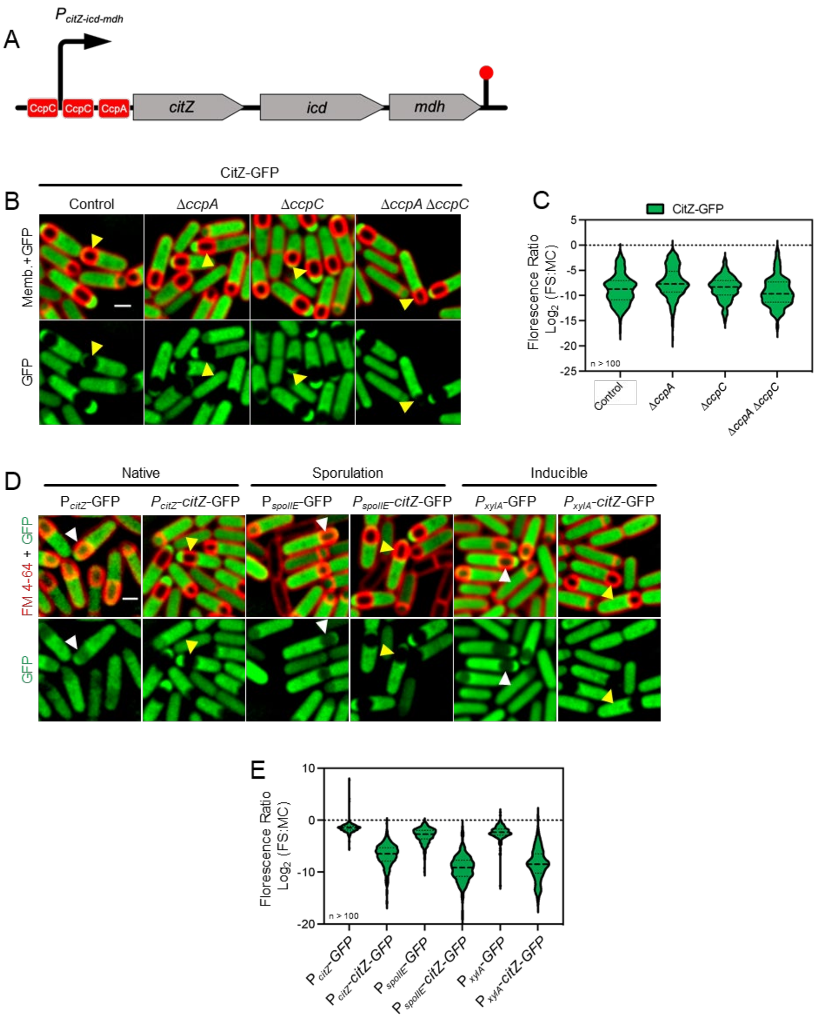


**Figure S3. CitZ-GFP depletion in the forespore occurs independently of transcriptional regulation.** (A) Diagram of the *citZ-icd-mdh* operon. The promoter (black arrow, *PcitZ-icd-mdh*), operators (CcpA and CcpC binding sites, red boxes), coding sequences (grey boxes) and transcription terminator (stick with red dot) are illustrated. (B) Visualization of CitZ-GFP (green) in sporulating cells lacking CcpA (Δ*ccpA*), CcpC (Δ*ccpC*) or both CcpA and CcpP (Δ*ccpA* Δ*ccpC*). Yellow arrowheads point to representative forespores. Membranes are stained with FM 4-64 (red). Scale bar, 1 µm. (C) Violin plots representing the $Log_2$ of the forespore-to-mother-cell (FS:MC) CitZ-GFP fluorescence ratio in sporulating cells from the strains shown in (B). Dotted lines within each violin plot delimit the middle 75% of the data and dashed lines indicate the median. A $Log_2$ value of 0 indicates an equal concentration of CitZ-GFP in the forespore and in the mother cell, and is marked with a dashed line. More than one hundred cells were analyzed for each strain. (D) Visualization of GFP fluorescent signal when GFP is produced alone (GFP) or fuse to CitZ (CitZ-GFP) from the following promoters: *citZ* promoter ($P_{citZ}$, native), the promoter of the sporulation gene *spoIIE* ($P_{spoIIE}$, sporulation) or the xylose inducible promoter of the *xylA* gene ($P_{xylA}$, inducible). Yellow and white arrowheads point to representative forespores without and with detectable GFP signal, respectively. Membranes are stained with FM 4-64 (red). Scale bar, 1 µm. (E) Violin plots representing the $Log_2$ of the FS:MC CitZ-GFP fluorescence ratio in sporulating cells from the strains shown in (D). See description of panel (C) for details about the violin plots. More than one hundred cells were analyzed for each strain.

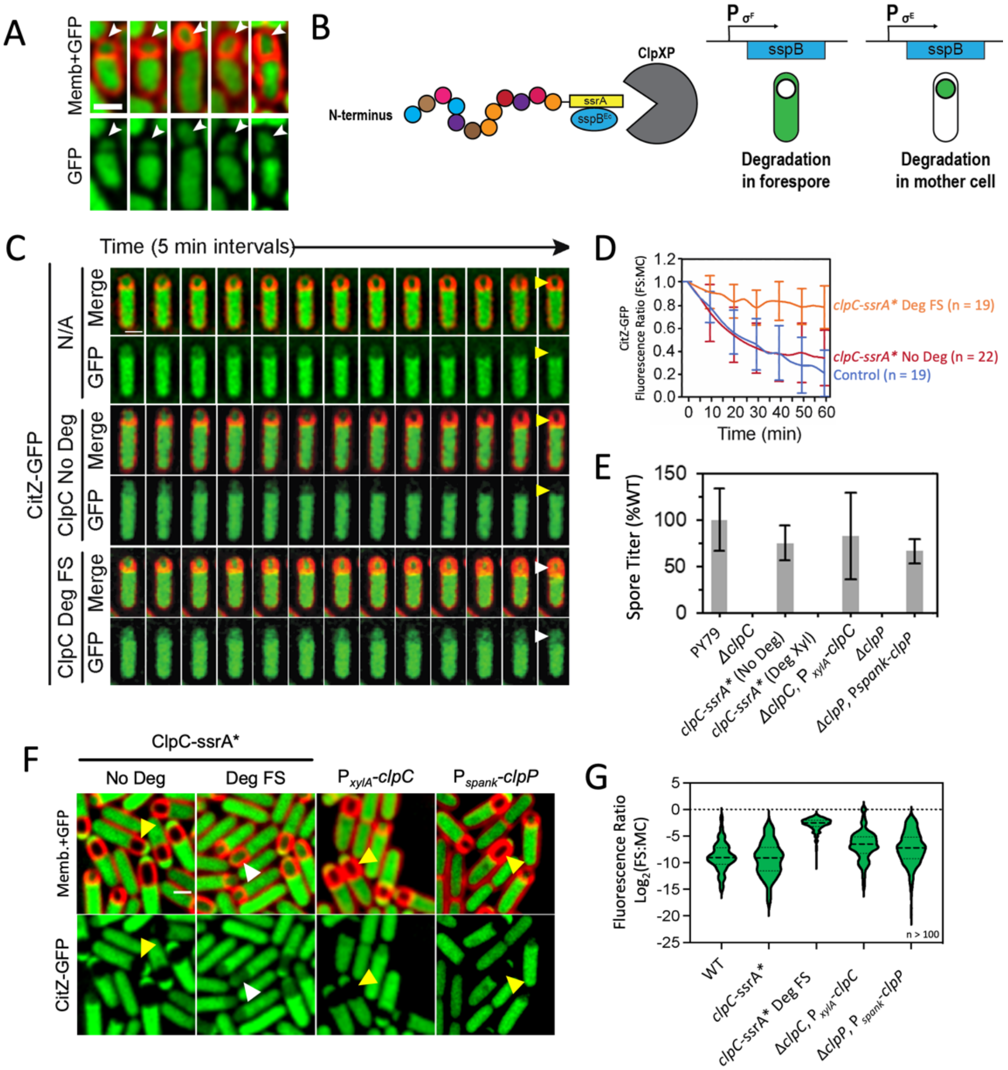


**Figure S4. ClpCP-mediated depletion of CitZ-GFP in the forespore is not related to indirect effects of ClpCP on $\sigma^F$ activation or to increased production of ClpCP in the forespore.** (A) Visualization of CitZ-GFP (green) in ClpP⁻ sporulating cells. Membranes are red. To allow sporulation initiation of the ClpP⁻ strain and be able to observe some sporulating cells, we used a strain that also lacked the RapA phosphatase, which partially rescued the sporulation initiation defect (Nanamiya et al. 2000). Every panel is a different sporulating cells. Forespores are on top, indicated by white arrowheads. Scale bar, 1 μm. (B) Diagram of Spatiotemporally Regulated Proteolysis (STRP). (C) Time-lapse fluorescence microscopy of sporulating cells producing CitZ-GFP (green) on a wild-type background (N/A, top two panels), in a strain in which ClpC is tagged with ssrA* but not degraded (ClpC No Deg, middle panels), and in a strain in which ClpC-ssrA* is degraded in the forespore (ClpC Deg FS). Forespores are at the top. Yellow and white arrowheads point to forespores in which CitZ-GFP is depleted and not depleted, respectively. Membranes are red (merge). Scale bar, 1 μm. (D) Quantification of the forespore-to-mother-cell (FS:MC) CitZ-GFP ratio of sporulating cells from the timelapses shown in (C) over time, relativized to the initial signal at $t_0$. Blue, wild-type background (WT); red, ClpC-ssrA* no degradation background (ClpC-ssrA* No Deg); orange, ClpC-ssrA* degraded in the forespore (ClpC-ssrA* Deg FS). Values represent the mean ± SD. (E) Spore titers of the following strains: wild type (WT), Δ*clpC*, ClpC-ssrA* no degradation (ClpC-ssrA* No Deg) strain, a strain in which ClpC-ssrA* is degraded by expressing *sspB^Ec^* from a xylose-inducible promoter (ClpC-ssrA* Deg Xyl), a complemented ΔclpC strain in which *clpC* is expressed from a xylose-inducible promoter (Δ*clpC*, $P_{xylA}$-*clpC*), ΔclpP, and a complemented Δ*clpP* strain in which *clpP* is expressed from an IPTG-inducible promoter (Δ*clpP*, $P_{spank}$-*clpP*). Values represent the mean ± SD of three independent experiments. (F) Visualization of CitZ-GFP in sporulating cells of the following strains: a strain in which ClpC is tagged with ssrA* but not degraded (first column), a strain in which ClpC is tagged with ssrA* and degraded in the forespore (second column), a strain in which *clpC* is expressed from a xylose-inducible promoter ($P_{xylA}$-*clpC*), and a strain in whch *clpP* is expressed from an IPTG-inducible promoter ($P_{spank}$-*clpP*). Membranes are in red. Yellow and white arrowheads indicate representative forespores lacking and containing CitZ-GFP, respectively. (G) Violin plots representing the $Log_2$ of the forespore-to-mother-cell CitZ-GFP fluorescence ratio in sporulating cells from the strains shown in (F). Dotted lines within each violin plot delimit the middle 75% of the data and dashed lines indicate the median. More than one hundred cells were analyzed for each strain. A $Log_2$ value of 0 indicates an equal concentration of CitZ-GP in the forespore and in the mother cell, and is marked with a dashed line.

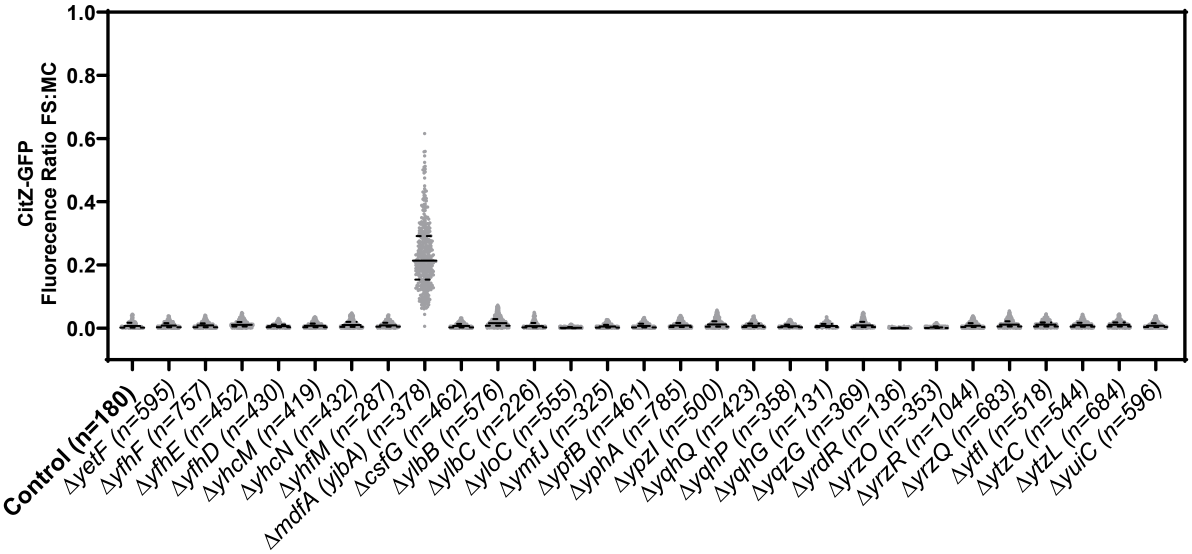


**Figure S5. Candidate-gene approach to identify *ori*-distal, $\sigma^F$-regulated genes required for CitZ-GFP depletion in the forespore.** Plots showing the ratios of the mean CitZ-GFP fluorescence in the forespore to the mean CitZ-GFP fluorescence in the mother cell (FS:MC) in sporulating cells of mutants containing deletions of the indicated genes. Every grey dot represents the CitZ-GFP fluorescent ratio of an individual cell, the dashed lines delimit the middle 75% of the data and the black solid lines indicate the median.

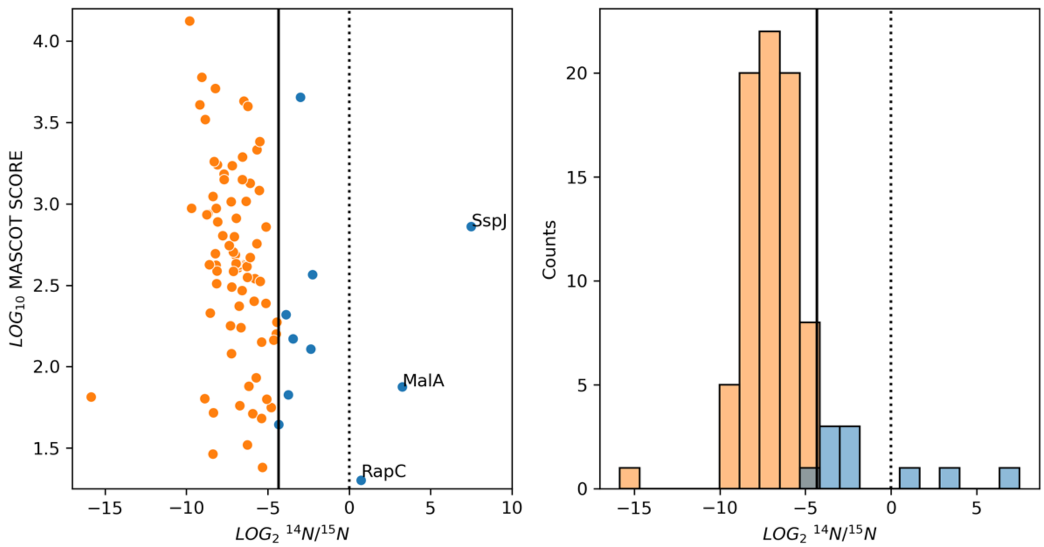


**Figure S6. Comparison of MdfA proteomic hits with previously published proteomic data.** Swarge *et al.* (Swarge et al. 2020) used an elegant approach to compared the relative levels of specific proteins in spores and vegetative cells using proteomics. The authors labelled proteins from exponentially-growing vegetative cells with $^{15}N$, and mixed them with proteins extracted from an equivalent number of spores that contained the most abundant nitrogen isotope, $^{14}N$. We checked if proteins enriched in MdfA$^-$ spores identified by us are more abundant in exponentially-growing vegetive cells than in dormant spores according to Swarge et al. (A) MASCOT scores versus $^{14}N/^{15}N$ protein isotopic ratio for individual proteins enriched in MdfA$^-$ spores according to our proteomics results. The MASCOT score indicates the combined spore and vegetative cell abundances of a protein. The $^{14}N/^{15}N$ protein isotopic ratio represents the relative levels of the protein in the spores compared to the vegetative cells. The vertical dotted and solid lines marks $^{14}N/^{15}N$ values of 1 and of 0.05, respectively. In principle, proteins with $^{14}N/^{15}N$ ratios below 1 are more abundant in exponentially-growing vegetative cells than in dormant spores, and proteins with $^{14}N/^{15}N$ ratios above 1 are more abundant in spores than in vegetative cells. Swarge *et al.* used a conservative $^{14}N/^{15}N$ cutoff value of 0.05 to classify proteins as predominantly present in vegetative cells (orange dots) or not (blue dots). Eighty-six of our proteomics hits were also identified by Swarge *et al.*, of which 83 had $^{14}N/^{15}N$ values below 1, and 76 below 0.05. Three of the MdfA$^-$ targets showed $^{14}N/^{15}N$ values above one, suggesting that they are more abundant in spores than in exponentially growing vegetative cells. One of them is the spore-specific protein SspJ, which is absent in vegetative cells (Bagyan et al. 1998). The other two, RapC and MalA, are expressed at higher levels in sporulating cultures than in exponentially-growing cells (Nicolas et al. 2012). The differential expression of these proteins during exponential growth and sporulation could therefore explain why these proteins are enriched in spores, in spite of being targeted by MdfA for degradation. (B) Histogram showing the distribution of $^{14}N/^{15}N$ values among the 86 proteins analyzed in (A). The orange and blue bars represent proteins that are predominantly found in vegetative cells and those that are not, respectively, according to the $^{14}N/^{15}N$ cutoff value of 0.05 used by Swarge *et al.* The vertical dotted and solid lines mark $^{14}N/^{15}N$ values of 1 and of 0.05, respectively.

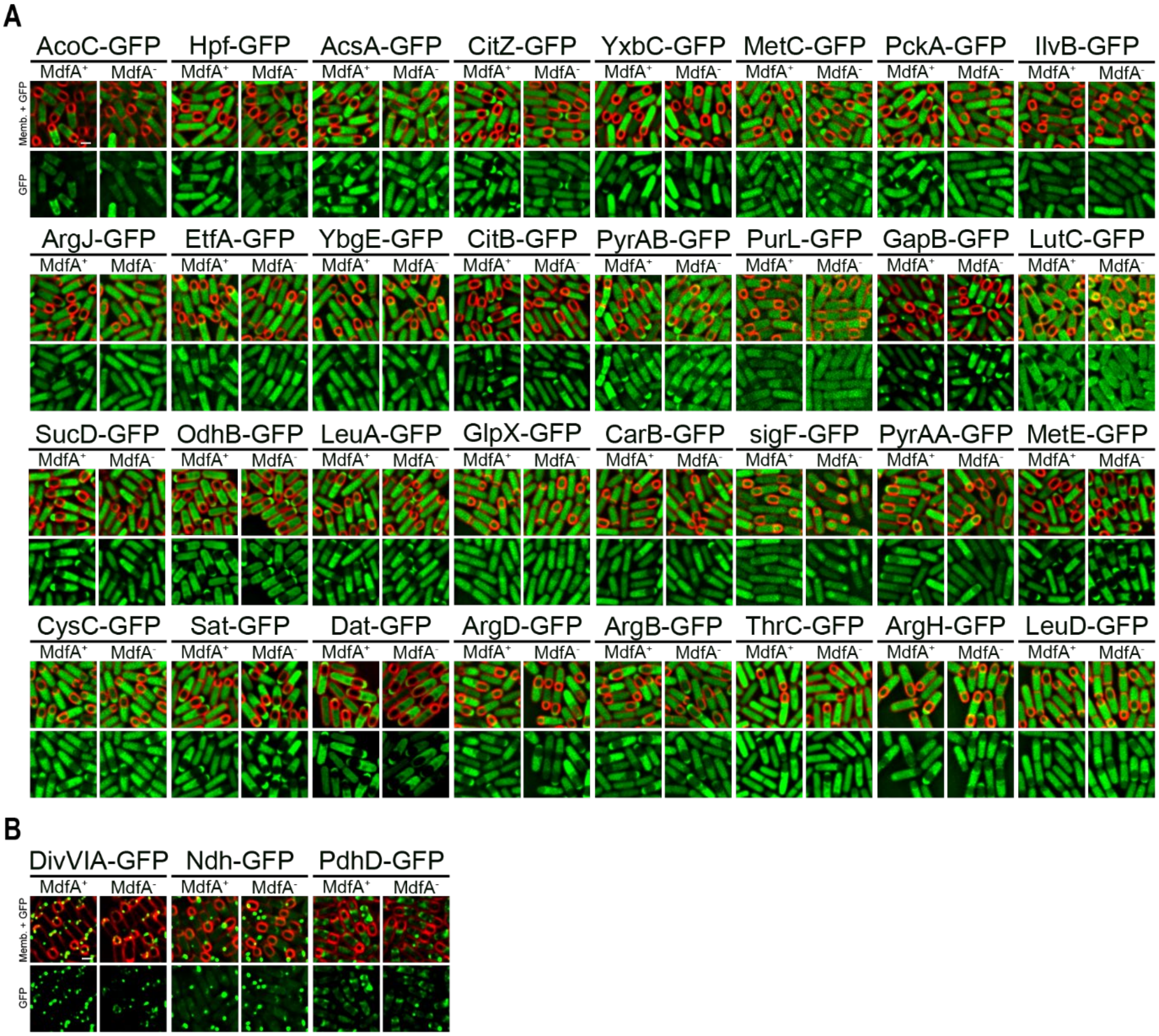


**Figure S7. Validation of putative MdfA targets identified by proteomics.** (A) GFP was fused to proteins enriched in MdfA$^-$ spores identified using proteomics. GFP signal from each fusion protein was visualized in MdfA$^+$ (wt) and MdfA$^-$ (Δ*mdfA*) sporulating cells. Membranes are stained with FM 4-64 (red). GFP in green. Scale is 1 μm. (B) Fusion proteins that form clusters were excluded from quantification. See Figure 5 for quantifications of the fusion proteins in (A).

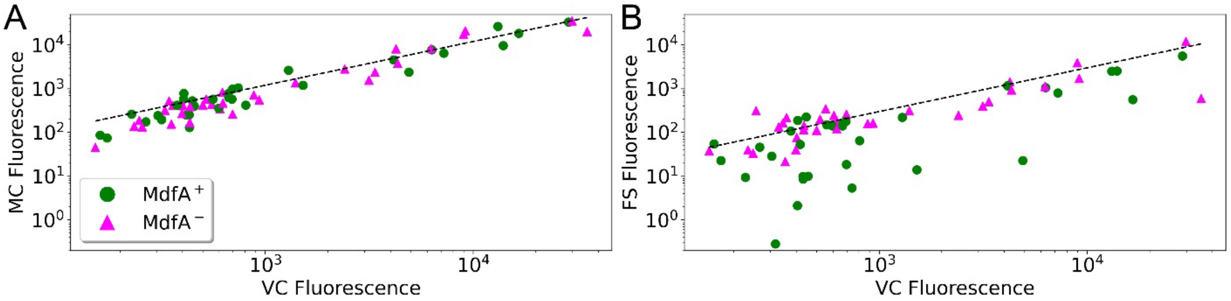


**Figure S8. MdfA mediates the depletion of proteins specifically in the forespore.** Scatter plots representing median GFP fluorescence intensity of mother cells (MC, A) or forespores (FS, B) versus that of vegetative cells (VC) present in the same sporulating cultures, for strains carrying the GFP fusions shown in Fig. S7. Every green circle and magenta triangle represent a different GFP fusion, in MdfA$^+$ and MdfA$^-$ backgrounds, respectively. The dotted lines represent the mother cell to vegetative cell (A) and the forespore to vegetative cell (B) fluorescence ratios expected if vegetative proteins were exclusively diluted in the forespore and concentrated in the mother cell due to forespore growth (see Fig. S2 for details). More than 150 forespores and mother cells, and more than 100 vegetative cells were analyzed for every strain. The only exception was ThrC-GFP, for which 23 and 44 forespores and mother cells were analyzed in MdfA$^+$ and MdfA$^-$ backgrounds, respectively.

**Table S1. Strains used in this study**

| Strain | Genotype or description | Reference, source or construction | Figure location |
|---|---|---|---|
| 1S138 | Δ*csfG::kan* | BGSC | |
| BER0269 | *coaD-ssrAΩkan* | pER14 → PY79 (Km) | |
| BER0389 | *coaD-ssrAΩerm* | pER32 → PY79 (MLS) | |
| BER0501 | *amyE::Pspank-clpPΩspec* | pER105 → PY79 (Sp) | |
| BER0502 | Δ*clpP::kan* | pER108 → PY79 (Km) | S4 |
| BER0504 | Δ*clpX::tet* | pER106→PY79 (Tc) | |
| BER0573 | *lacA::PspoIIE-gfpΩerm* | pER138 → PY79 (MLS) | S1 |
| BER0707 | Δ*clpC::tet* | pER161→PY79 (Tc) | S4 |
| BER0715 | Δ*clpX::tet, citZ-gfpΩkan* | BER0504→JLG2113 (Tc) | 3 |
| BER0716 | Δ*clpC::tet, citZ-gfpΩkan* | BER0707→JLG2113 (Tc) | 3 |
| BER0717 | Δ*lon::cat, citZ-gfpΩkan* | KP419→JLG2113 (Cm) | 3 |
| BER0834 | Δ*ccpC::tet* | pER178 → PY79 (Tc) | |
| BER0835 | *clpC-ssrA*Ωkan* | pER179→PY79 (Km) | |
| BER0889 | *clpC-ssrA*Ωkan, ΔxylA::sspB*$^{Ec}$Ω*cat* | BER0835→JLG364 (Km) | S4 |
| BER0905 | *citZ-gfp* | pCrePA→JLG2113 (Em) | |
| BER0906 | *icd-gfp* | pCrePA→JLG2114 (Em) | |
| BER0920 | *clpC-ssrA*Ωkan, citZ-gfp* | BER0835→BER0905 (Km) | |
| BER0961 | *citB-gfpΩkan* | (Riley et al. 2021) | 5, S7, S8 |
| BER1091 | *guaB-gfpΩkan* | (Riley et al. 2021) | 1 |
| BER1095 | *rpoZ-gfpΩkan* | (Riley et al. 2021) | 1 |
| BER1098 | *metE-gfpΩkan* | (Riley et al. 2021) | 1 |
| BER1101 | *leuD-gfpΩkan* | (Riley et al. 2021) | 1 |
| BER1102 | *argH-gfpΩkan* | (Riley et al. 2021) | 1 |
| BER1107 | *gapB-gfpΩkan* | (Riley et al. 2021) | 5, S7, S8 |
| BER1215 | *clpC-ssrA*Ωkan, amyE::P*$_{spoIIQ}$-*sspB*$^{Ec}$*Ωcat* | BER0835→JLG170 (Km) | S4 |
| BER1220 | *citZ-gfp*, Δ*sigF::kan* | BER0606 → BER0905 (Km) | 3 |
| BER1221 | *citZ-fp*, Δ*sigE::erm* | BER0605 → BER0905 (Km) | 3 |
| BER1222 | Δ*spoIIQ::spc*, *citZ-gfp* | KP575→BER905 (Sp) | 2 |
| BER1228 | Δ*spoIIQ::spc*, *icd-gfp* | KP575→BER906 (Sp) | 2 |
| BER1232 | *amyE::P*$_{spoIIQ}$-*sspB*$^{Ec}$*Ωcat, clpC-ssrA*Ωkan, citZ-gfp* | JLG170→BER0920 (Cm) | S4 |
| BER1375 | *citZ-gfpΩkan, spoIIIEATP- (G467S; ATPase mutant)* | JLG2113→KP541 (Km) | 3 |
| BER1385 | Δ*clpE::cat* | pER324→PY79 (Cm) | |

| | | | |
|---|---|---|---|
| BER1386 | Δ*clpQY::kan* | pER325→PY79 (Km) | |
| BER1387 | Δ*ispA::erm* | pER326→PY79 (Em) | |
| BER1395 | Δ*ispA::erm, citZ-gfpΩkan* | BER1387→JLG2113 (MLS) | 3 |
| BER1396 | Δ*clpE::cat, citZ-gfpΩkan* | BER1385→JLG2113 (Cm) | 3 |
| BER1426 | Δ*clpQY* | pCrePA→BER1386 (Em) | |
| BER1428 | *citZ-gfpΩkan,* Δ*clpQY* | JLG2113→BER1426 (Km) | 3 |
| BER1431 | Δ*yetF::erm, citZ-gfpΩkan* | BKE07140→JLG2113 (Em) | 3, S5 |
| BER1432 | Δ*yfhF::erm, citZ-gfpΩkan* | BKE08510→JLG2113 (Em) | 3, S5 |
| BER1433 | Δ*yfhE::erm, citZ-gfpΩkan* | BKE08500→JLG2113 (Em) | 3, S5 |
| BER1434 | Δ*yfhD::erm, citZ-gfpΩkan* | BKE08490→JLG2113 (Em) | 3, S5 |
| BER1435 | Δ*yhcM::erm, citZ-gfpΩkan* | BKE09140→JLG2113 (Em) | 3, S5 |
| BER1436 | Δ*yhcN::erm, citZ-gfpΩkan* | BKE09150→JLG2113 (Em) | 3, S5 |
| BER1437 | Δ*yhfM::erm, citZ-gfpΩkan* | BKE10280→JLG2113 (Em) | 3, S5 |
| BER1438 | Δ*yjbA::erm, citZ-gfpΩkan* | BKE11410→JLG2113 (Em) | 3, 5, S5, S7, S8 |
| BER1439 | Δ*csfG::kan, citZ-gfpΩkan* | 1S138→JLG2113 (Em) | 3, S5 |
| BER1440 | Δ*ylbB::erm, citZ-gfpΩkan* | BKE14950→JLG2113 (Em) | 3, S5 |
| BER1441 | Δ*ylbC::erm, citZ-gfpΩkan* | BKE14960→JLG2113 (Em) | 3, S5 |
| BER1442 | Δ*yloC::erm, citZ-gfpΩkan* | BKE15660→JLG2113 (Em) | 3, S5 |
| BER1443 | Δ*ymfJ::erm, citZ-gfpΩkan* | BKE16880→JLG2113 (Em) | 3, S5 |
| BER1444 | Δ*ypfB::erm, citZ-gfpΩkan* | BKE22900→JLG2113 (Em) | 3, S5 |
| BER1445 | Δ*yphA::erm, citZ-gfpΩkan* | BKE22860→JLG2113 (Em) | 3, S5 |
| BER1446 | Δ*ypzI::erm, citZ-gfpΩkan* | BKE22869→JLG2113 (Em) | 3, S5 |
| BER1447 | Δ*yqhQ::erm, citZ-gfpΩkan* | BKE24490→JLG2113 (Em) | 3, S5 |
| BER1448 | Δ*yqhP::erm, citZ-gfpΩkan* | BKE24500→JLG2113 (Em) | 3, S5 |
| BER1449 | Δ*yqhG::erm, citZ-gfpΩkan* | BKE24590→JLG2113 (Em) | 3, S5 |
| BER1450 | Δ*yqzG::erm, citZ-gfpΩkan* | BKE24650→JLG2113 (Em) | 3, S5 |
| BER1451 | Δ*yrdR::erm, citZ-gfpΩkan* | BKE26620→JLG2113 (Em) | 3, S5 |
| BER1452 | Δ*yrzO::erm, citZ-gfpΩkan* | BKE26619→JLG2113 (Em) | 3, S5 |
| BER1453 | Δ*yrzR::erm, citZ-gfpΩkan* | BKE27469→JLG2113 (Em) | 3, S5 |
| BER1454 | Δ*yrzQ::erm, citZ-gfpΩkan* | BKE27468→JLG2113 (Em) | 3, S5 |
| BER1455 | Δ*ytfI::erm, citZ-gfpΩkan* | BKE29510→JLG2113 (Em) | 3, S5 |
| BER1456 | Δ*ytzC::erm, citZ-gfpΩkan* | BKE30470→JLG2113 (Em) | 3, S5 |
| BER1457 | Δ*ytzL::erm, citZ-gfpΩkan* | BKE30739→JLG2113 (Em) | 3, S5 |
| BER1458 | Δ*yuiC::erm, citZ-gfpΩkan* | BKE32070→JLG2113 (Em) | 3, S5 |
| BER1461 | Δ*clpC::tet,* Δ*yjbA::erm, citZ-gfpΩkan* | BER0707→BER1438 (Tc) | |
| BER1465 | Δ*yjbA::erm, gapB-gfpΩkan* | BER1479 → BER1107 (MLS | 5, S7, S8 |
| BER1479 | Δ*yjbA::erm* | BKE11410 → PY79 (MLS) | 4 |

| | | | |
|---|---|---|---|
| BER1486 | *yjbA-gfpΩkan* | pER344→PY79 (Km) | 3 |
| BER1492 | Δ*yjbA::erm, citB-gfpΩkan* | BER1479 → BER0961(MLS) | 5, S7, S8 |
| BER1493 | Δ*yjbA::erm, odhB-gfpΩkan* | BER1479 → JLG2107 (MLS) | 5, S7, S8 |
| BER1494 | Δ*yjbA::erm, metE-gfpΩkan* | BER1098 → BER1479 (Km) | 5, S7 |
| BER1506 | *xylA::clpCΩcat,* Δ*clpC::tet* | pER342→BER0707 (Cm) | |
| BER1519 | *citZ-gfpΩkan, xylA::clpCΩcat,* Δ*clpC::tet* | JLG2113→BER1506 (Km) | |
| BER1541 | Δ*ctpA::tet* | pER354→PY79 (Tc) | |
| BER1542 | Δ*ctpB::tet* | pER355→PY79 (Tc) | |
| BER1543 | Δ*ftsH::tet* | pER356→PY79 (Tc) | |
| BER1544 | Δ*gpr::tet* | pER357→PY79 (Tc) | |
| BER1551 | Δ*ctpA::tet, citZ-gfpΩkan* | BER1541→JLG2113 (Tc) | 3 |
| BER1552 | Δ*ctpB::tet, citZ-gfpΩkan* | BER1542→JLG2113 (Tc) | 3 |
| BER1553 | Δ*ftsH::tet, citZ-gfpΩkan* | BER1543→JLG2113 (Tc) | 3 |
| BER1554 | Δ*gpr::tet, citZ-gfpΩkan* | BER1544→JLG2113 (Tc) | 3 |
| BER1576 | *metE-gfpΩkan,* Δ*spoIIQ::spc* | BER1098→KP575 (Km) | S3 |
| BER1577 | *prs-gfpΩkan,* Δ*spoIIQ::spc* | BER1110→KP575 (Km) | S3 |
| BER1578 | *rpoZ-gfpΩkan,* Δ*spoIIQ::spc* | BER1095→KP575 (Km) | S3 |
| BER1604 | Δ*yjbA::erm, sucD-gfpΩkan* | JLG2111 → BER1479 (Km) | 5, S7, S8 |
| BER1637 | $P_{citZ}$-*gfpΩkan* | pER375→PY79 (Km) | |
| BER1640 | $P_{citZ}$-*gfpΩkan,* Δ*spoIIQ::spc* | BER1637→KP575 (Km) | 2 |
| BER1686 | *leuA-gfpΩkan* | pER394 → PY79 (Km) | 5, S7, S8 |
| BER1691 | Δ*yjbA::erm, leuA-gfpΩkan* | BER1686 → BER1479 (Km) | 5, S7, S8 |
| BER1710 | *carB-gfpΩkan* | pER397 → PY79 (Km) | 5, S7, S8 |
| BER1711 | *ilvB-gfpΩkan* | pER398 → PY79 (Km) | 5, S7, S8 |
| BER1715 | *pckA-gfpΩkan* | pER403 → PY79 (Km) | 5, S7, S8 |
| BER1718 | *pdhD-gfpΩkan* | pER406 → PY79 (Km) | 5, S7, S8 |
| BER1735 | *ilvB-gfpΩkan,* Δ*yjbA::erm* | BER1711 → BER1479 (Km) | 5, S7, S8 |
| BER1737 | *pckA-gfpΩkan,* Δ*yjbA::erm* | BER1715 → BER1479 (Km) | 5, S7, S8 |
| BER1738 | *pdhD-gfpΩkan,* Δ*yjbA::erm* | BER1718 → BER1479 (Km) | 5, S7, S8 |
| BER1753 | Δ*yjbA::erm, carB-gfpΩkan* | BER1710 → BER1479 (Km) | 5, S7, S8 |
| BER1764 | Δ*yjbA::erm, argH-gfpΩkan* | BER1102 → BER1479 (Km) | 5, S7, S8 |
| BER1800 | *etfA-gfpΩkan* | BJAL003 → PY79 (Km) | 5, S7, S8 |
| BER1801 | Δ*yjbA::erm, etfA-gfpΩkan* | BJAL003 → BER1479 (Km) | 5, S7, S8 |
| BER1818 | *ndh-gfpΩkan* | pER422 → PY79 (Km) | 5, S7, S8 |
| BER1819 | *sigF-gfpΩkan* | pER423 → PY79 (Km) | 5, S7, S8 |
| BER1820 | Δ*yjbA::erm, ndh-gfpΩkan* | BER1818 → BER1479 (Km) | 5, S7, S8 |
| BER1821 | Δ*yjbA::erm, sigF-gfpΩkan* | BER1819 → BER1479 (Km) | 5, S7, S8 |

| | | | |
|---|---|---|---|
| BER1859 | *ywrK::amyE::PspoIIQ-lacZ (cat), ΔyjbA::erm, amyE::PyjbA-yjbA(E116K) (kan)* | (Massoni et al. 2024) | |
| BER1861 | *glpX-gfpΩkan* | pER424 → PY79 (Km) | 5, S7, S8 |
| BER1862 | *acoC-gfpΩkan* | pER425 → PY79 (Km) | 5, S7, S8 |
| BER1863 | *lutC-gfpΩkan* | pER426 → PY79 (Km) | 5, S7, S8 |
| BER1864 | *acsA-gfpΩkan* | pER427 → PY79 (Km) | 5, S7, S8 |
| BER1865 | *ΔyjbA::erm, glpX-gfpΩkan* | BER1861 → BER1479 (Km) | 5, S7, S8 |
| BER1866 | *ΔyjbA::erm, lutC-gfpΩkan* | BER1863 → BER1479 (Km) | 5, S7, S8 |
| BER1867 | *ΔyjbA::erm, acoC-gfpΩkan* | BER1862 → BER1479 (Km) | 5, S7, S8 |
| BER1868 | *ΔyjbA::erm, acsA-gfpΩkan* | BER1864 → BER1479 (Km) | 5, S7, S8 |
| BER1876 | *pyrAA-gfpΩkan* | pER430 → PY79 (Km) | 5, S7, S8 |
| BER1877 | *pyrAB-gfpΩkan* | pER431 → PY79 (Km) | 5, S7, S8 |
| BER1881 | *clpC(K85E)Ωkan* | pER435 → PY79 (Km) | |
| BER1885 | *yjbA(E116K)Ωerm* | pER437 → PY79 (MLS) | |
| BER1886 | *citZ-gfp , clpC(K85E)Ωkan* | BER1881 → BER0905 (Km) | |
| BER1889 | *ΔyjbA::erm, pyrAA-gfpΩkan* | BER1876 → BER1479 (Km) | 5, S7, S8 |
| BER1890 | *ΔyjbA::erm, pyrAB-gfpΩkan* | BER1877 → BER1479 (Km) | 5, S7, S8 |
| BER1896 | *citZ-gfp , clpC(WT)Ωkan* | BER1897 → BER0905 (Km) | |
| BER1897 | *clpCΩkan* | pER315 → PY79 (Km) | |
| BER1905 | *citZ-gfp, clpC(WT)Ωkan, yjbA(E116K)Ωerm* | BER1885 → BER1896 (MLS) | 3 |
| BER1907 | *citZ-gfp, clpC(K85E)Ωkan, yjbA(E116K)Ωerm* | BER1885 → BER1886 (MLS) | 3 |
| BER1914 | *argJ-gfpΩkan* | pER442 → PY79 (Km) | 5, S7, S8 |
| BER1915 | *argB-gfpΩkan* | pER443 → PY79 (Km) | 5, S7, S8 |
| BER1916 | *argD-gfpΩkan* | pER444 → PY79 (Km) | 5, S7, S8 |
| BER1920 | *ΔyjbA::erm, argJ-gfpΩkan* | BER1914 → BER1479 (Km) | 5, S7, S8 |
| BER1921 | *ΔyjbA::erm, argB-gfpΩkan* | BER1915 → BER1479 (Km) | 5, S7, S8 |
| BER1922 | *ΔyjbA::erm, argD-gfpΩkan* | BER1916 → BER1479 (Km) | 5, S7, S8 |
| BER1927 | *ybgE-gfpΩkan* | pER441 → PY79 (Km) | 5, S7, S8 |
| BER1928 | *ΔyjbA::erm, ybgE-gfpΩkan* | BER1927 → BER1479 (Km) | 5, S7, S8 |
| BER1929 | *ΔyjbA::erm, leuD-gfpΩkan* | BER1101 → BER1479 (Km) | 5, S7, S8 |
| BER1933 | *yxbC-gfpΩkan* | pER447 → PY79 (Km) | 5, S7, S8 |
| BER1940 | *ΔccpC::tet, citZ-gfp* | BER0834 → BER0905 (Tc) | S1 |
| BER1943 | *ΔyjbA::erm, yxbC-gfpΩkan* | BER1933 → BER1479 (Km) | 5, S7, S8 |
| BER1946 | *purL-gfpΩkan* | pER452 → PY79 (Km) | 5, S7, S8 |
| BER1948 | *divIVA-gfpΩkan* | pER454 → PY79 (Km) | 5, S7, S8 |
| BER1949 | *thrC-gfpΩkan* | pER455 → PY79 (Km) | 5, S7, S8 |
| BER1950 | *hpf-gfpΩkan* | pER456 → PY79 (Km) | 5, S7, S8 |
| BER1952 | *ΔccpA::erm* | pER458 → PY79 (Km) | |

| | | | |
|---|---|---|---|
| BER1955 | *ΔyjbA::erm, purL-gfpΩkan* | BER1946 → BER1479 (Km) | 5, S7, S8 |
| BER1957 | *ΔyjbA::erm, divIVA-gfpΩkan* | BER1948 → BER1479 (Km) | 5, S7, S8 |
| BER1958 | *ΔyjbA::erm, thrC-gfpΩkan* | BER1949 → BER1479 (Km) | 5, S7, S8 |
| BER1959 | *ΔyjbA::erm, hpf-gfpΩkan* | BER1950 → BER1479 (Km) | 5, S7, S8 |
| BER1963 | *ΔccpA::erm, citZ-gfp* | BER1952 → BER1479 (Km) | S1 |
| BER1966 | *metC-gfpΩkan* | pER461 → PY79 (Km) | 5, S7, S8 |
| BER1967 | *cysC-gfpΩkan* | pER462 → PY79 (Km) | 5, S7, S8 |
| BER1968 | *sat-gfpΩkan* | pER463 → PY79 (Km) | 5, S7, S8 |
| BER1969 | *lacA::PspoIIE-citZ-gfpΩerm* | pER464 → PY79 (Km) | S1 |
| BER1970 | *xylA::gfpΩcat* | pER465 → PY79 (Cm) | S1 |
| BER1972 | *ΔyjbA::erm, metC-gfpΩkan* | BER1966 → BER1479 (Km) | 5, S7, S8 |
| BER1973 | *ΔyjbA::erm, cysC-gfpΩkan* | BER1967 → BER1479 (Km) | 5, S7, S8 |
| BER1974 | *ΔyjbA::erm, sat-gfpΩkan* | BER1968 → BER1479 (Km) | 5, S7, S8 |
| BER1982 | *ΔclpP::kan, citZ-gfp* | BER0502 → BER0905 (Km) | 3 |
| BER1984 | *ΔccpA::erm, ΔccpC::tet, citZ-gfp* | BER1952 → BER1940 (MLS) | S1 |
| BER1985 | *amyE::Pspank-clpPΩspec, citZ-gfp* | BER0501 → BER0905 (Sp) | S4 |
| BER1986 | *ΔclpP::kan, amyE::Pspank-clpPΩspec, citZ-gfp* | BER0502 → BER1985 (Km) | S4 |
| BER1989 | *xylA::citZ-gfpΩcat* | pER467 → PY79 (Cm) | S1 |
| BKE07140 | *trpC2 ΔyetF::erm* | (Koo et al. 2017) | |
| BKE08490 | *trpC2 ΔyfhD::erm* | (Koo et al. 2017) | |
| BKE08500 | *trpC2 ΔyfhE::erm* | (Koo et al. 2017) | |
| BKE08510 | *trpC2 ΔyfhF::erm* | (Koo et al. 2017) | |
| BKE09140 | *trpC2 ΔyhcM::erm* | (Koo et al. 2017) | |
| BKE09150 | *trpC2 ΔyhcN::erm* | (Koo et al. 2017) | |
| BKE10280 | *trpC2 ΔyhfM::erm* | (Koo et al. 2017) | |
| BKE12430 | *trpC2 ΔrapA::erm* | (Koo et al. 2017) | |
| BKE14950 | *trpC2 ΔylbB::erm* | (Koo et al. 2017) | |
| BKE14960 | *trpC2 ΔylbC::erm* | (Koo et al. 2017) | |
| BKE15660 | *trpC2 ΔyloC::erm* | (Koo et al. 2017) | |
| BKE16880 | *trpC2 ΔymfJ::erm* | (Koo et al. 2017) | |
| BKE22860 | *trpC2 ΔyphA::erm* | (Koo et al. 2017) | |
| BKE22869 | *trpC2 ΔypzI::erm* | (Koo et al. 2017) | |
| BKE22900 | *trpC2 ΔypfB::erm* | (Koo et al. 2017) | |
| BKE24490 | *trpC2 ΔyqhQ::erm* | (Koo et al. 2017) | |
| BKE24500 | *trpC2 ΔyqhP::erm* | (Koo et al. 2017) | |
| BKE24590 | *trpC2 ΔyqhG::erm* | (Koo et al. 2017) | |
| BKE24650 | *trpC2 ΔyqzG::erm* | (Koo et al. 2017) | |
| BKE26619 | *trpC2 ΔyrzO::erm* | (Koo et al. 2017) | |
| BKE26620 | *trpC2 ΔyrdR::erm* | (Koo et al. 2017) | |

| | | | |
|---|---|---|---|
| BKE27468 | *trpC2* Δ*yrzQ::erm* | (Koo et al. 2017) | |
| BKE27469 | *trpC2* Δ*yrzR::erm* | (Koo et al. 2017) | |
| BKE29510 | *trpC2* Δ*ytfI::erm* | (Koo et al. 2017) | |
| BKE30470 | *trpC2* Δ*ytzC::erm* | (Koo et al. 2017) | |
| BKE30739 | *trpC2* Δ*ytzL::erm* | (Koo et al. 2017) | |
| BKE32070 | *trpC2* Δ*yuiC::erm* | (Koo et al. 2017) | |
| BKE34540 | *trpC2* Δ*clpP::erm* | (Koo et al. 2017) | |
| BKK11410 | *trpC2* Δ*yjbA::kan* | (Koo et al. 2017) | |
| BJAL003 | *etfA-GFPΩkan* | pJAL011→ PY79 | |
| JLG0170 | *amyE::P$_{spoIIQ}$-sspB$^{Ec}$Ωcat* | (Shin et al. 2015) | |
| JLG0364 | *ΔxylA::sspB$^{Ec}$Ωcat* | (Lamsa et al. 2016) | |
| JLG0626 | *ΔspoIIQ::erm* | (Ojkic et al. 2016) | |
| JLG0963 | *amyE::PsspE(2G)-sspB$^{Ec}$Ωcat* | (Lopez-Garrido et al. 2018) | |
| JLG2107 | *odhB-gfpΩkan* | (Riley et al. 2021) | 5, S7, S8 |
| JLG2111 | *sucD-gfp* | (Riley et al. 2021) | 5, S7, S8 |
| JLG2113 | *citZ-gfpΩkan* | (Riley et al. 2021) | 1, 2, 3, 5, S2, S3, S4, S5, S7, S8 |
| JLG2114 | *icd-gfpΩkan* | (Riley et al. 2021) | 1, 2, 5, S7, S8 |
| JLG5876 | Δ*rapA::erm* | BKE12430 → PY79 (MLS) | |
| JLG5979 | *dat-gfpΩerm* | pJLG1214 → PY79 (MLS) | 5, S7, S8 |
| JLG6008 | Δ*rapA* | pDR244 → JLG5876 (Sp) | |
| JLG6060 | *dat-gfpΩerm* Δ*yjbA::kan* | BKK11410 → JLG5979 (Km) | 5, S7, S8 |
| JLG6214 | Δ*rapA citZ-gfpΩkan* | JLG2113 → JLG6008 (Km) | |
| JLG6267 | Δ*clpP::erm* Δ*rapA citZ-GFPΩkan* | BKE34540 → JLG6214 (MLS) | S4 |
| KP0161 | Δ*sigE::erm* | KP strain collection | |
| KP0324 | Δ*sigF::kan* | KP strain collection | |
| KP0419 | Δ*lon::cat* | (Schmidt et al. 1994) | |
| KP0541 | *spoIIIEATP- (G467S; ATPase mutant)* | (Sharp and Pogliano 1999) | |
| KP0575 | Δ*spoIIQ::spc* | (Londoño-Vallejo et al. 1997) | |
| PY79 | Wild type | (Youngman et al. 1984) | 4, S4 |

[a] Plasmid or genomic DNA used (right side the arrow) to transform an existing strain (left side the arrow) to create a new strain are listed.

| Table S2. Plasmids used in this study | |
|---|---|
| **Plasmid** | **Description/Reference** |
| pCrePA | (Pomerantsev et al. 2006) |
| pDG1662 | (Guérout-Fleury et al. 1996) |
| pDG1731 | (Guérout-Fleury et al. 1996) |
| pDR110 | A gift from David Rudner |
| pDR244 | (Meeske et al. 2015) |
| pER14 | *coaD-ssrA*Ωkan* |
| pER32 | *coaD-ssrA*Ωerm* |
| pER105 | *amyE::Pspank-rbs-clpPΩspec* |
| pER106 | Δ*clpC::tet* |
| pER108 | Δ*clpC::kan* |
| pER128 | (Riley et al. 2018) |
| pER138 | *lacA::PspoIIE-gfpΩerm* |
| pER161 | Δ*clpC::tet* |
| pER178 | Δ*ccpC::tet* |
| pER179 | *clpC-ssrA*Ωkan* |
| pER315 | *clpCΩkan* |
| pER324 | Δ*clpE::cat* |
| pER325 | Δ*clpQY::kan* |
| pER326 | Δ*ispA::erm* |
| pER342 | *xylA::clpCΩcat* |
| pER344 | *yjbA-gfpΩkan* |
| pER354 | Δ*ctpA::tet* |
| pER355 | Δ*ctpB::tet* |
| pER356 | Δ*ftsH::tet* |
| pER357 | Δ*gpr::tet* |
| pER360 | *xylA::clpC-gfpΩcat* |
| pER375 | $P_{citZ}$*-gfpΩkan* |
| pER394 | *leuA-gfpΩkan* |
| pER397 | *carB-gfpΩkan* |
| pER398 | *ilvB-gfpΩkan* |
| pER403 | *pckA-gfpΩkan* |
| pER406 | *pdhD-gfpΩkan* |
| pER422 | *ndh-gfpΩkan* |
| pER423 | *sigF-gfpΩkan* |
| pER424 | *glpX-gfpΩkan* |
| pER425 | *acoC-gfpΩkan* |
| pER426 | *lutC-gfpΩkan* |
| pER427 | *acsA-gfpΩkan* |

| | |
|---|---|
| pER430 | *pyrAA-gfpΩkan* |
| pER431 | *pyrAB-gfpΩkan* |
| pER435 | *clpC (K85E)Ωkan* |
| pER437 | *yjbA(E116K)Ωerm* |
| pER441 | *ybgE-gfpΩkan* |
| pER442 | *argJ-gfpΩkan* |
| pER443 | *argB-gfpΩkan* |
| pER444 | *argD-gfpΩkan* |
| pER447 | *yxbC-gfpΩkan* |
| pER452 | *purL-gfpΩkan* |
| pER454 | *divIVA-gfpΩkan* |
| pER455 | *thrC-gfpΩkan* |
| pER456 | *hpf-gfpΩkan* |
| pER458 | Δ*ccpA::erm* |
| pER461 | *metC-gfpΩkan* |
| pER462 | *cysC-gfpΩkan* |
| pER463 | *sat-gfpΩkan* |
| pER464 | *lacA::PspoIIE-citZ-gfpΩerm* |
| pER465 | *xylA::gfpΩcat* |
| pER467 | *xylA::citZ-gfpΩcat* |
| pJAL011 | *etfA-gfpΩkan* |
| pJLG3 | (Shin et al. 2015) |
| pJLG38 | (Shin et al. 2015) |
| pJLG67 | (Lamsa et al. 2016) |
| pJLG396 | *rpsB-gfpΩerm* |
| pJLG1214 | *dat-gfpΩerm* |
| pWH1520 | (Rygus and Hillen 1991) |

| Table S3. Primers used in this study | |
|---|---|
| **Primer** | **Squence**[a] |
| oER52 | gggttaacgcgtaatccatgGACTTATTCGCAGGGAGCGG |
| oER53 | catcatttgctgcgctagcTCCTTGTCTGAATTTTTGCTGAAGC |
| oER54 | cactggagttgtcccaattGGGTTTTGACTCATCCTTTTTTTG |
| oER55 | cacatttccccgaaaagtgcGGCTTCGTTCTTGCTATGGG |
| oER417 | ttctgctccctcgctcaggcggccgcTTTTCGCTACGCTCAAATCC |
| oER418 | cagggagcactggtcaacgctagcGACTGTAAAAAGTACAGTCGGC |
| oER419 | ttctgctccctcgctcaggcggccgcTCCTTTAACTCTGGCAACCC |
| oER420 | cagggagcactggtcaacgctagcATGTGATAACTCGGCGTATG |
| oER421 | ttctgctccctcgctcaggcggccgcATGAGAGAGGAAGAAAACGG |
| oER421 | ttctgctccctcgctcaggcggccgcATGAGAGAGGAAGAAAACGG |
| oER422 | cagggagcactggtcaacgctagcAATTGGGACAACTCCAGTG |
| oER425 | ttctgctccctcgctcaggcggccgcTCCTTTAACTCTGGCAACCC |
| oER425 | ttctgctccctcgctcaggcggccgcTCCTTTAACTCTGGCAACCC |
| oER426 | cagggagcactggtcaacgctagcATGTGATAACTCGGCGTATG |
| oER455 | gggttaacgcgtaatccatgATAGAGAAGAACTGGGGGAGC |
| oER456 | gcctgagcgagggagcagaaGCCTGATATGCAAAGTCGTC |
| oER457 | gcgttgaccagtgctccctgTAACACAACCTGCAAGAGCTG |
| oER458 | cacatttccccgaaaagtgcAAATGCGGAGAAACAAGAGC |
| oER459 | gggttaacgcgtaatccatgCCTGACTCTTGAGGCGATTG |
| oER460 | gcctgagcgagggagcagaaGTGTTTTTCCACAGAACGAGC |
| oER461 | gcgttgaccagtgctccctgAAAGACGGCACTGAGGTAAGC |
| oER462 | cacatttccccgaaaagtgcATCGAGTTCCACAAAGACAGC |
| oER463 | acgacgaagCTTACCTAAAAGGTGAAGGAGGAG |
| oER464 | agcagcgcatgcTTACTTTTTGTCTTCTGTGTGAGTC |
| oER718 | gggttaacgcgtaatccatgCTTTTGGCGGCTGTTTGC |
| oER719 | gcctgagcgagggagcagaaAGCAGCAATGCCCTCTCCTTC |
| oER720 | gcgttgaccagtgctccctgCGGTGCCCGTCCGTTAAG |
| oER721 | cacatttccccgaaaagtgcCCTGTGACACACTGCCTGGAG |
| oER728 | ATGGAACATGCAGAACACGG |
| oER732 | cacatttccccgaaaagtgcGTGAGTCCAGTTGTAGAGCGG |
| oER735 | cactggagttgtcccaattTCCTTTTAATGAATCTGGCATTG |
| oER736 | gcgcttgcgctgctagcGTAAGCCAGGCTGAAAAGTCC |
| oER775 | gggttaacgcgtaatccatgCAAGTTCTTGAAGACGGACG |
| oER776 | catcatttgctgcgctagcATTCGTTTTAGCAGTCGTTTTTACG |
| oER777 | cactggagttgtcccaattTATAGAAGACGGAAATGAGGC |
| oER778 | cacatttccccgaaaagtgcGTGACGGATTATTAATGCCG |
| oER779 | gcgcttgcgctgctagcATTCGTTTTAGCAGTCGTTTTTACG |

| | |
|---|---|
| oER789 | gggttaacgcgtaatccatgAAGTAGCGCACGTGCAAGTC |
| oER790 | gcctgagcgagggagcagaaGAAGCTGCATGTCGTCCTC |
| oER791 | gcgttgaccagtgctccctgTGAAACGTGCAAACAGATG |
| oER792 | cacatttccccgaaaagtgcCATGCAAGAAAACCTCTCG |
| oER1104 | TCACAGTGGCAATCTCCCC |
| oER1105 | TCTTTCTGCTTTTTCCATTCCTC |
| oER1132 | ccgttttcttcctctctcatTTAATTCGTTTTAGCAGTCGTTTTTACG |
| oER1156 | gggttaacgcgtaatccatgGATATGACTTGGCATTGATTCG |
| oER1157 | gcctgagcgagggagcagaaCAATGTTGACAACGCATTTGC |
| oER1158 | gcgttgaccagtgctccctgCGAGCAAAATAACAATCAGCG |
| oER1159 | cacatttccccgaaaagtgcTAACGACTGCTGAGATTGTCC |
| oER1162 | gggttaacgcgtaatccatgCTGTTAGAGCTGCTCTATGC |
| oER1163 | gcctgagcgagggagcagaaAAGATGACATAAAGGGCCTCC |
| oER1164 | gcgttgaccagtgctccctgGAAAAGCTCGGAACGATAGC |
| oER1165 | cacatttccccgaaaagtgcCCATTCCGACAACTGTTGC |
| oER1168 | gggttaacgcgtaatccatgCTCCTGTTTCTCATAAACTCC |
| oER1169 | gcctgagcgagggagcagaaCTGCTCATTCGTCACATACG |
| oER1170 | gcgttgaccagtgctccctgGCAATCACATTTGTTGACCC |
| oER1171 | cacatttccccgaaaagtgcTTCCAATACGCTTCAAGTGC |
| oER1190 | aagcttaaggaggtatacatATGATGTTTGGAAGATTTACAG |
| oER1191 | ggatttgagcgtagcgaaaaTTAATTCGTTTTAGCAGTCG |
| oER1198 | gggttaacgcgtaatccatgTGTTTTGTCGTAACAGACGG |
| oER1199 | gcgcttgcgctgctagcGTTGACCTTTGATTCGTTTTCC |
| oER1200 | cactggagttgtcccaattAAAAAACCGCTCTTTGCAAAG |
| oER1201 | cacatttccccgaaaagtgcCAACGACATCTTTACGATTCC |
| oER1203 | CATATATAACATCTCCTTTTCAATAAATTTCC |
| oER1235 | gggttaacgcgtaatccatgCTCTTGAAATTTCTTGAAGAGAGG |
| oER1236 | gcctgagcgagggagcagaaTTAAACACCACCTTTTTCTTCTTAC |
| oER1237 | gcgttgaccagtgctccctgAAAAAACCATACGCGGCTG |
| oER1238 | cacatttccccgaaaagtgcCCTTGTTCCATTATGGCAATCG |
| oER1241 | gggttaacgcgtaatccatgCCGTTTGTTTAAAGTGGTCG |
| oER1242 | gcctgagcgagggagcagaaCCTTTACTCCTCCTGTATGC |
| oER1243 | gcgttgaccagtgctccctgAGGAGATGGTATTTCATCTCCTG |
| oER1244 | cacatttccccgaaaagtgcGACCAAATATCTGCTGAACAGC |
| oER1247 | gggttaacgcgtaatccatgACATTTCCGTTGGCAATCG |
| oER1248 | gcctgagcgagggagcagaaTCCTTACCTCCTCCCACAG |
| oER1249 | gcgttgaccagtgctccctgTTCGCTTTCTTTCTAAAAAAAC |
| oER1250 | cacatttccccgaaaagtgcGCATTTACGATTCTGTCTGC |
| oER1253 | gggttaacgcgtaatccatgATCCCCATTATTGAGGATCG |

| | |
|---|---|
| oER1254 | gcctgagcgagggagcagaaCTCACTCTTTTTCATACAGGG |
| oER1255 | gcgttgaccagtgctccctgACAGTTCTACTTCCTCTAGC |
| oER1256 | cacatttccccgaaaagtgcCTCTTTCTTCCTCCATCACC |
| oER1266 | ggatttgagcgtagcgaaaaTCATTTATACAGTTCATCCATGCCATG |
| oER1316 | GAAGAACTGTTTACAGGCG |
| oER1341 | gggttaacgcgtaatccatgGCTGACTGCTTTAGGATTCC |
| oER1342 | gcgcttgcgctgctagcTGATCCGACAGCTGTGTG |
| oER1343 | cactggagttgtcccaattAAGAAAGGAGAACGGTTAACTTG |
| oER1344 | cacatttccccgaaaagtgcCTGATCACTCGTTCAATTTCCG |
| oER1383 | gggttaacgcgtaatccatgTGTCGTCAGCAAAGTAATGG |
| oER1384 | gcgcttgcgctgctagcCTGTGTGCATGATGCCAC |
| oER1385 | cactggagttgtcccaattATGCACACAGTGACGCAAAC |
| oER1386 | cacatttccccgaaaagtgcCCACGTTATTTCCGTCACCG |
| oER1389 | gggttaacgcgtaatccatgGTAGGCCAGCATCAAATGTGG |
| oER1390 | gcgcttgcgctgctagcAGGTTTCACCCCCACCATTTC |
| oER1391 | cactggagttgtcccaattTTGAAAAGAATTATCACATTGACTGTG |
| oER1392 | cacatttccccgaaaagtgcCTTTTACCATTTCAATCTCTCCC |
| oER1401 | gggttaacgcgtaatccatgGCAAACTAACGAAGGAGCAGG |
| oER1403 | cactggagttgtcccaattAAAACAAAAGCCAAGAGC |
| oER1404 | cacatttccccgaaaagtgcCGTGCTGTTAATCCATGG |
| oER1405 | gcgcttgcgctgctagcTACGAGAGGGCCGCCTG |
| oER1415 | gggttaacgcgtaatccatgGGTATCGAAATGACGGACC |
| oER1417 | cactggagttgtcccaattTTTTCATATCAAAAACAGCCCC |
| oER1418 | cacatttccccgaaaagtgcCTTTCATCCAAGAATGCAGC |
| oER1419 | gcgcttgcgctgctagcTTTTACGATGTGAATCGGAC |
| oER1435 | gcgcttgcgctgctagcGCCATCCGTATGATCCATTTG |
| oER1454 | gggttaacgcgtaatccatgAACAAGGACGTTGAGGACG |
| oER1455 | gcgcttgcgctgctagcTGGACGGATTACAAGATTTGG |
| oER1456 | cactggagttgtcccaattCAAACAGAAAAAGCGGGTG |
| oER1457 | cacatttccccgaaaagtgcGCATTCGCTTTCAGAATGG |
| oER1460 | gggttaacgcgtaatccatgAACAGCTTTCACCAACTGC |
| oER1461 | gcgcttgcgctgctagcTAAAATTAATGCTGCGGGTTC |
| oER1462 | cactggagttgtcccaattGAAAAGCAGGTGAAAACGAC |
| oER1463 | cacatttccccgaaaagtgcCTAGTGAAGACGGAATCTCG |
| oER1466 | gggttaacgcgtaatccatgCTGATTGAAACGGATTCAACCG |
| oER1467 | gcgcttgcgctgctagcGCGGTCAGAGACGAGAATATATG |
| oER1468 | cactggagttgtcccaattACTCAGGAAGCCCGGCAG |
| oER1469 | cacatttccccgaaaagtgcTGAATGAAGTTCGAGTATGCAGCG |
| oER1472 | gggttaacgcgtaatccatgAATCAAGGCAACGAGCTACC |

| | |
|---|---|
| oER1473 | gcgcttgcgctgctagcATCCTCCATTGTTGACAGATCTC |
| oER1474 | cactggagttgtcccaattAAGGCGGCGTTGACACAG |
| oER1475 | cacatttccccgaaaagtgcCGCTTGGATCACCGATGAGG |
| oER1496 | gggttaacgcgtaatccatgAACGGAAATGTGACGTCATCG |
| oER1497 | gcgcttgcgctgctagcCGCGTTTTGGCATACCGC |
| oER1498 | cactggagttgtcccaattATGCCAAAACGCGTAGAC |
| oER1499 | cacatttccccgaaaagtgcCTGTTCCGCCTAATGTATATGC |
| oER1502 | gggttaacgcgtaatccatgGCTGAGAAGAGTGGATATTACG |
| oER1503 | gcgcttgcgctgctagcTATAGTGACTGCCGCCTC |
| oER1504 | cactggagttgtcccaattATGAAAAAAGCGTATTTGACAG |
| oER1505 | cacatttccccgaaaagtgcTCACATCTGTGACAAATCCG |
| oER1517 | cacatttccccgaaaagtgcCGAGTGTTCCGATCATTCC |
| oER1520 | CATTATACTCCTAGAGCTgAAAAAGTCATTGAGCTTTC |
| oER1522 | gcctgagcgagggagcagaatcaaaaaaccgctctttgcaaagagcggtttttttcaGTTGACCTTTGATTCGTTTTCC |
| oER1523 | gcgttgaccagtgctccctgAAAAAACCGCTCTTTGCAAAG |
| oER1526 | gggttaacgcgtaatccatgATGTTATTTCTTCATGATGTGTG |
| oER1533 | gggttaacgcgtaatccatgACATCCACGAACGACATGG |
| oER1534 | gcgcttgcgctgctagcCGTGCGATAGCTCGCG |
| oER1535 | cactggagttgtcccaattTAAAGGGGAGCGAAATGAAG |
| oER1536 | cacatttccccgaaaagtgcTTCATGATTCCATCGACATCC |
| oER1539 | gggttaacgcgtaatccatgTTTCGTTGCTGAGCTTGC |
| oER1540 | gcgcttgcgctgctagcTGAAACAGCCTCCTTTGC |
| oER1541 | cactggagttgtcccaattAAAGGAGGCTGTTTCATGAG |
| oER1542 | cacatttccccgaaaagtgcAAGCAGAAGGATCATTGTATGG |
| oER1551 | gggttaacgcgtaatccatgTCGAAGATGAGTATGTGAGGG |
| oER1552 | gcgcttgcgctgctagcCACTTCCACTGTCCAGTTAAAC |
| oER1553 | cactggagttgtcccaattAAATCGAAAAAGAACCTGCC |
| oER1554 | cacatttccccgaaaagtgcGACATTTGCAAAGCTGACG |
| oER1557 | gggttaacgcgtaatccatgGTCGTTTTGCAAGGAAAAGC |
| oER1558 | gcgcttgcgctgctagcCTGGTTTACAGCGGAATGATG |
| oER1559 | cactggagttgtcccaattTTTTTTTTCGATATAAAGGCATAAAAATTC |
| oER1560 | cacatttccccgaaaagtgcTGAGAGCGATATGTTTATTCCC |
| oER1565 | gggttaacgcgtaatccatgATTCAAGGTGAGAAAACATGG |
| oER1566 | gcgcttgcgctgctagcGAACTTATAACTTGTCAAAAGCTG |
| oER1567 | cactggagttgtcccaattTAAATACTAGATAAGGGGATTAATGAC |
| oER1568 | cacatttccccgaaaagtgcCTGCCAGTAAAGGAATTTCG |
| oER1589 | gggttaacgcgtaatccatgCAGCATTTCAAACAAGCAGG |
| oER1590 | gcgcttgcgctgctagcAGCCTTTGATTTCAGCAAGC |

| | |
|---|---|
| oER1591 | cactggagttgtcccaattGTATGGAAAGGAGCTATCCC |
| oER1592 | cacatttccccgaaaagtgcGCGACAATCATTTCTGTTTCG |
| oER1607 | gggttaacgcgtaatccatgCAAATGATATTCACAACAAGACG |
| oER1608 | gcgcttgcgctgctagcTTCCTTTTCCTCAAATACAGC |
| oER1609 | cactggagttgtcccaattATTCTCTGATTATCTTGACATTTTC |
| oER1610 | cacatttccccgaaaagtgcCGGTAGATATCTTCTTCCTCC |
| oER1613 | gggttaacgcgtaatccatgGGAAGCGCCTGATGTTTTAGCG |
| oER1614 | gcgcttgcgctgctagcTACACGGGCCGCTCCTTTTAC |
| oER1615 | cactggagttgtcccaattAGCATCCTTGAATATGTAAAAGG |
| oER1616 | cacatttccccgaaaagtgcAAGGAATGACAACGACTACG |
| oER1619 | gggttaacgcgtaatccatgTCATGTCGAGAGGAAGATCG |
| oER1620 | gcgcttgcgctgctagcTTCAGTCGGTTCAATTAAGCC |
| oER1621 | cactggagttgtcccaattTGAAGAGAAGCCTTCCGTG |
| oER1622 | cacatttccccgaaaagtgcAGACGTTAATGTTTTCCCTTCC |
| oER1631 | gggttaacgcgtaatccatgAAACAGCAACGAGAAACACG |
| oER1632 | gcctgagcgagggagcagaaCCTAAAACCACTCCTTTTACTGG |
| oER1633 | gcgttgaccagtgctccctgGAAAAACAAAGAGCAAGCTTC |
| oER1659 | gggttaacgcgtaatccatgTGGGAAGCAGCTGTATAAGC |
| oER1660 | gcgcttgcgctgctagcACGAACCGAAACTGGAGC |
| oER1661 | cactggagttgtcccaattAAAAGAAGCCGGGGATAGTC |
| oER1662 | cacatttccccgaaaagtgcAGCGAAAATCGAGATTTTTTCG |
| oER1665 | gggttaacgcgtaatccatgGCATAAAAGCTCCATTCTCTGG |
| oER1666 | gcgcttgcgctgctagcGCTCAATTTTTGTTTAACAAACTCG |
| oER1667 | cactggagttgtcccaattATATGATCTGCTGCGTTCTTC |
| oER1668 | cacatttccccgaaaagtgcAGCGCTTTCCACTTTTCC |
| oER1671 | gggttaacgcgtaatccatgTTGTTCTTGAATCCGTTAGTAGG |
| oER1672 | gcgcttgcgctgctagcAGATACGCCTACTTCTTCTTTTTC |
| oER1673 | cactggagttgtcccaattGGAGGACTGGCAGTGAC |
| oER1674 | cacatttccccgaaaagtgcTCCTGTAAAGAACGGAATCTCG |
| oJLG0007 | AATTGGGACAACTCCAGTG |
| oJLG0077 | GCTAGCAGCGCAAGCGC |
| oJLG0095 | CATGGATTACGCGTTAACCC |
| oJLG0096 | GCACTTTTCGGGGAAATGTG |
| oJLG0097 | ATGAGAGAGGAAGAAAACGG |
| oJLG0184 | GCTAGCGCAGCAAATGATG |
| oJLG0187 | TTTTCGCTACGCTCAAATCC |
| oJLG0188 | *cccttcagtaaagctgcggt*GACTGTAAAAAGTACAGTCGGC |
| oJLG0253 | TCCTTTAACTCTGGCAACCC |
| oJLG0254 | ATGTGATAACTCGGCGTATG |

| | |
|---|---|
| oJLG0550 | ATGTATACCTCCTTAAGCTTAATTGTTATC |
| oJLG0860 | catacgccgagttatcacatAACTTTTCACTGGAGTTGTCCC |
| oJLG0861 | gggttgccagagttaaaggaCTCTCATTTATCCTCCAAGTGC |
| oJLG0885 | cacatttccccgaaaagtgcGTCTTGCTCTGTCACAGCTG |
| oJLG0886 | cttgcgcttgcgctgctagcTACGTACCACAATTTTGTTCCG |
| oJLG1218 | gggttaacgcgtaatccatgTGCTCTTTGCTGTATCCGTC |
| oJLG1406 | TACCTAGATTTAGATGTCTAAAAAGC |
| oJLG1407 | TTATTATTTCCTTCCTCTTTTCTAC |
| oJLG1408 | gtagaaaagaggaaggaaataataaATGAACGAGAAAAATATAAAACACAG |
| oJLG1409 | gctttttagacatctaaatctaggtaGCAGTTTATGCATCCCTTAAC |
| oJLG1696 | catggattacgcgttaacccaggtctagaggatcgatctg |
| oJLG1699 | gctagcagcgcaagcgcaGCTAGCGCAGCAAATGATG |
| oJLG3036 | gggttaacgcgtaatccatgCTACTGCCTATACGTTTACG |
| oJLG3037 | tgcgcttgcgctgctagcTGAAATGCTAGCAGCCTGT |
| oJLG3038 | cactggagttgtcccaattATGAACGAAGAAAAGAGCCC |
| oJLG3039 | cacatttccccgaaaagtgcCAACTTTCGAAACTGACAATG |

[a]In capital letters are shown the regions of the primer that anneals to the template. Homology regions for Gibson assembly and restriction sites are shown in lowercase.

### Supplemental Methods

#### Details of plasmid construction

**pER14**. This plasmid was constructed by assembling the following four fragments by Gibson Assembly (New England Biolabs): (i) 3' region of *coaD* amplified with primers oER52 and oER53 from genomic DNA of *B. subtilis* PY79; (ii) ssrA-kan fragment amplified from pJLG3 with primers oJLG184 and oJLG7; (iii) *coaD* downstream region amplified from genomic DNA of PY79 with primers oER54 and oER55; (iv) a DNA fragment encompassing the spectinomycin resistance gene, the origin of replication, and the ampicillin resistance gene from pDG1662 (Guérout-Fleury et al. 1996), amplified with primers oJLG95 and oJLG96.

**pER32**. Constructed by two-part Gibson Assembly reaction: pER014 was amplified with primers oJLG860 and oJLG861 to exclude *kan* cassette, and assembled with an *erm* cassette amplified from genomic DNA of JLG626 with primers oJLG253 and oJLG254.

**pER105.** The coding sequence of clpP, including the native ribosome-binding site, was amplified from genomic DNA of PY79 using oER463 and oER464. The fragment was digested with HindIII and SphI and ligated into pDR110 cut with the same enzymes.

**pER106**. This plasmid was constructed by assembling the following four fragments by Gibson Assembly (New England Biolabs): (i) a *clpX* upstream homology fragment amplified with primers oER459 and oER460 from genomic DNA of *B. subtilis* PY79; (ii) a tetracycline resistance gene amplified with primers oJLG885 and oJLG886 from pWH1520 (Rygus and Hillen 1991), and then re-amplified with primers oER425 and oER426; (iii) a *clpX* downstream homology fragment amplified with primers oER461 and oER462 from genomic DNA of *B. subtilis* PY79; and (iv) a DNA fragment encompassing the spectinomycin resistance gene, the origin of replication, and the ampicillin resistance gene from pDG1662 (Guérout-Fleury et al. 1996), amplified with primers oJLG95 and oJLG96.

**pER108.** This plasmid was constructed by assembling the following four fragments by Gibson Assembly (New England Biolabs): (i) a *clpP* upstream homology fragment amplified with primers oER455 and oER456 from genomic DNA of *B. subtilis* PY79; (ii) a kanamycin resistance gene amplified with primers oER421 and oER422 from BER0269; (iii) a *clpP* downstream homology fragment amplified with primers oER457 and oER458 from genomic DNA of *B. subtilis* PY79; and (iv) a DNA fragment encompassing the spectinomycin resistance gene, the origin of replication, and the ampicillin resistance gene from pDG1662 (Guérout-Fleury et al. 1996), amplified with primers oJLG95 and oJLG96.

**pER138.** This plasmid was constructed by assembling the following 2 fragments by Gibson Assembly (New England Biolabs): (i) a 5706 bp fragment of *lacA::PspoIIEΩerm* amplified from pER128 (Riley et al 2018) using primers oER1104 and oER1105; (ii) *gfpΩkan* fragment amplified with primers oJLG7 and oJLG77 from pJLG38 (Shin et al. 2015).

**pER161**. This plasmid was constructed by assembling the following four fragments by Gibson Assembly (New England Biolabs): (i) a fragment of 905 bp amplified with primers oER718 and oER719 from genomic DNA of *B. subtilis* PY79, including the first 117 nucleotides of *clpC* coding sequence and the 788 nucleotides immediately upstream of it; (ii) a tetracycline resistance gene amplified with primers oJLG885 and oJLG886 from pWH1520 (Rygus and Hillen 1991), and then re-amplified with primers oER425 and oER426; (iii) a fragment of 825 bp encompassing the last 154 nucleotides of *clpC* coding sequence and the region immediately downstream of it, amplified with primers oER720 and oER721 from genomic DNA of *B. subtilis* PY79; and (iv) a DNA fragment encompassing the spectinomycin resistance gene, the origin of replication, and the ampicillin resistance gene from pDG1662 (Guérout-Fleury et al. 1996), amplified with primers oJLG95 and oJLG96.

**pER178.** This plasmid was constructed by assembling the following four fragments by Gibson Assembly (New England Biolabs): (i) a *ccpA* upstream homology fragment amplified with primers oER789 and oER790 from genomic DNA of *B. subtilis* PY79; (ii) a tetracycline resistance gene amplified with primers oJLG885 and oJLG886 from pWH1520 (Rygus and Hillen 1991), and then re-amplified with primers oER425 and oER426; (iii) a *ccpA* downstream homology fragment amplified with primers oER791 and oER792 from genomic DNA of *B. subtilis* PY79; and (iv) a DNA fragment encompassing the spectinomycin resistance gene, the origin of replication, and the ampicillin resistance gene from pDG1662 (Guérout-Fleury et al. 1996), amplified with primers oJLG95 and oJLG96.

**pER179**. This plasmid was constructed by assembling the following four fragments by Gibson Assembly (New England Biolabs): (i) 3' region of *clpC* coding sequence (not including the stop codon) amplified with primers oER775 and oER776 from genomic DNA of *B. subtilis* PY79; (ii) *ssrA*Ωkan* fragment amplified with primers oJLG7 and oJLG184 from pJLG3 (Shin et al. 2015); (iii) region immediately downstream of the last amino acid-coding codon of *clpC*, amplified with primers oER777 and oER778 from genomic DNA of *B. subtilis* PY79; and (iv) a DNA fragment encompassing the spectinomycin resistant gene, the origin of replication, and the ampicillin resistant gene from pDG1662 (Guérout-Fleury et al. 1996), amplified with primers oJLG95 and oJLG96.

**pER315.** This plasmid was constructed by assembling the following three fragments by Gibson Assembly (New England Biolabs): (i) a *clpC* upstream homology fragment amplified with primers oER718 and oER1132 from genomic DNA of BER0835; (ii) a *clpC-kan* fragment amplified with primers oJLG97 and oER778 from genomic DNA of BER0835; and (iii) a DNA fragment encompassing the spectinomycin resistance gene, the origin of replication, and the ampicillin resistance gene from pDG1662 (Guérout-Fleury et al. 1996), amplified with primers oJLG95 and oJLG96.

**pER324**. This plasmid was constructed by assembling the following four fragments by Gibson Assembly (New England Biolabs): (i) a fragment of 595 bp amplified with primers oER1156 and oER1157 from genomic DNA of *B. subtilis* PY79, encompassing the region immediately upstream of the *clpE* coding sequence; (ii) a chloramphenicol resistance gene amplified with primers oJLG187 and oJLG188 from pDG1662 (Guérout-Fleury et al. 1996), and then re-amplified with primers oER417 and oER418; (iii) a fragment of 539 bp encompassing the last 12 nucleotides of *clpE* coding sequence and the region immediately downstream of it, amplified with primers oER1158 and oER1159 from genomic DNA of *B. subtilis* PY79; and (iv) a DNA fragment encompassing the spectinomycin resistance gene, the origin of replication, and the ampicillin resistance gene from pDG1662 (Guérout-Fleury et al. 1996), amplified with primers oJLG95 and oJLG96.

**pER325**. This plasmid was constructed by assembling the following four fragments by Gibson Assembly (New England Biolabs): (i) a fragment of 523 bp amplified with primers oER1162 and oER1163 from genomic DNA of *B. subtilis* PY79, including the first 10 nucleotides of *clpQY* coding sequence and the nucleotides immediately upstream of it; (ii) a kanamycin resistance gene amplified with primers oER421 and oER422 from pJLG3 (Shin et al. 2015); (iii) a fragment of 550 bp encompassing the last 54 nucleotides of *clpQY* coding sequence and the region immediately downstream of it, amplified with primers oER1164 and oER1165 from genomic DNA of *B. subtilis* PY79; and (iv) a DNA fragment encompassing the spectinomycin resistance gene, the origin of replication, and the ampicillin resistance gene from pDG1662 (Guérout-Fleury et al. 1996), amplified with primers oJLG95 and oJLG96.

**pER326**. This plasmid was constructed by assembling the following four fragments by Gibson Assembly (New England Biolabs): (i) a fragment of 572 bp amplified with primers oER1168 and oER1169 from genomic DNA of *B. subtilis* PY79, including the first 45 nucleotides of *ispA* coding sequence and the nucleotides immediately upstream of it; (ii) an erythromycin resistance gene amplified with primers oER419 and oER420 from pDG1731 (Guérout-Fleury et al. 1996); (iii) a fragment of 526 bp encompassing the last 25 nucleotides of *ispA* coding sequence and the region immediately downstream of it, amplified with primers oER1170 and oER1171 from genomic DNA of *B. subtilis* PY79; and (iv) a DNA fragment encompassing the spectinomycin resistance gene, the origin of replication, and the ampicillin resistance gene from pDG1662 (Guérout-Fleury et al. 1996), amplified with primers oJLG95 and oJLG96.

**pER342**. This plasmid was constructed by assembling the following two fragments by Gibson Assembly (New England Biolabs): (i) a fragment of 2433 bp amplified with primers oER1190 and oER1191 from genomic DNA of *B. subtilis* PY79, encompassing the entire *clpC* coding sequence, including the stop codon; (ii) a DNA fragment encompassing the chloramphenicol resistance gene, the origin of replication, the xylose repressor *xylR*, and the xylA promoter $P_{xylA}$ from pJLG67 (Lamsa et al. 2016), amplified with primers oJLG187 and oJLG550.

**pER344**. This plasmid was constructed by assembling the following four fragments by Gibson Assembly (New England Biolabs): (i) 3' region of *yjbA* coding sequence (not including the stop codon) amplified with primers oER1198 and oER1199 from genomic DNA of *B. subtilis* PY79; (ii) *gfpΩkan* fragment amplified with primers oJLG7 and oJLG77 from pJLG38 (Shin et al. 2015); (iii) region immediately downstream of the *yjbA* stop codon, amplified with primers oER1200 and oER1201 from genomic DNA of *B. subtilis* PY79; and (iv) a DNA fragment encompassing the spectinomycin resistance gene, the origin of replication, and the ampicillin resistance gene from pDG1662 (Guérout-Fleury et al. 1996), amplified with primers oJLG95 and oJLG96.

**pER354**. This plasmid was constructed by assembling the following four fragments by Gibson Assembly (New England Biolabs): (i) a fragment of 525 bp amplified with primers oER1235 and oER1236 from genomic DNA of *B. subtilis* PY79, including region immediately upstream of the *ctpA* coding sequence; (ii) a tetracycline resistance gene amplified with primers oJLG885 and oJLG886 from pWH1520 (Rygus and Hillen 1991), and then re-amplified with primers oER425 and oER426; (iii) a fragment of 517 bp encompassing the region immediately downstream of the *ctpA* coding sequence, amplified with primers oER1237 and oER1238 from genomic DNA of *B. subtilis* PY79; and (iv) a DNA fragment encompassing the spectinomycin resistance gene, the origin of replication, and the ampicillin resistance gene from pDG1662 (Guérout-Fleury et al. 1996), amplified with primers oJLG95 and oJLG96.

**pER355**. This plasmid was constructed by assembling the following four fragments by Gibson Assembly (New England Biolabs): (i) a fragment of 554 bp amplified with primers oER1241 and oER1242 from genomic DNA of *B. subtilis* PY79, including region immediately upstream of the *ctpB* coding sequence; (ii) a tetracycline resistance gene amplified with primers oJLG885 and oJLG886 from pWH1520 (Rygus and Hillen 1991), and then re-amplified with primers oER425 and oER426; (iii) a fragment of 519 bp encompassing the region immediately downstream of the *ctpB* coding sequence, amplified with primers oER1243 and oER1244 from genomic DNA of *B. subtilis* PY79; and (iv) a DNA fragment encompassing the spectinomycin resistance gene, the origin of replication, and the ampicillin resistance gene from pDG1662 (Guérout-Fleury et al. 1996), amplified with primers oJLG95 and oJLG96.

**pER356**. This plasmid was constructed by assembling the following four fragments by Gibson Assembly (New England Biolabs): (i) a fragment of 541 bp amplified with primers oER1247 and oER1248 from genomic DNA of *B. subtilis* PY79, including region immediately upstream of the *ftsH* coding sequence; (ii) a tetracycline resistance gene amplified with primers oJLG885 and oJLG886 from pWH1520 (Rygus and Hillen 1991), and then re-amplified with primers oER425 and oER426; (iii) a fragment of 535 bp encompassing the region immediately downstream of the *ftsH* coding sequence, amplified with primers oER1249 and oER1250 from genomic DNA of *B. subtilis* PY79; and (iv) a DNA fragment encompassing the spectinomycin resistance gene, the origin of replication, and the ampicillin resistance gene from pDG1662 (Guérout-Fleury et al. 1996), amplified with primers oJLG95 and oJLG96.

**pER357**. This plasmid was constructed by assembling the following four fragments by Gibson Assembly (New England Biolabs): (i) a fragment of 513 bp amplified with primers oER1253 and oER1254 from genomic DNA of *B. subtilis* PY79, including the first 15 nucleotides of the *gpr* coding sequence and the 498 nucleotides immediately upstream of it; (ii) a tetracycline resistance gene amplified with primers oJLG885 and oJLG886 from pWH1520 (Rygus and Hillen 1991), and then re-amplified with primers oER425 and oER426; (iii) a fragment of 516 bp encompassing the the region immediately downstream of the *gpr* coding sequence, amplified with primers oER1255 and oER1256 from genomic DNA of *B. subtilis* PY79; and (iv) a DNA fragment encompassing the spectinomycin resistance gene, the origin of replication, and the ampicillin resistance gene from pDG1662 (Guérout-Fleury et al. 1996), amplified with primers oJLG95 and oJLG96.

**pER360**. Constructed by two-piece Gibson assembly reaction: inverse PCR of pER342 was performed using oER779 and oJLG187, and the resulting fragment was assembled with *gfp* amplified using oJLG77 and oER1266.

**pER375**. This plasmid was constructed by assembling the following two fragments by Gibson Assembly (New England Biolabs): (i) a fragment of 731 bp amplified with primers oER1203 and oJLG1218 from genomic DNA of *B. subtilis* PY79, encompassing the *citZ* start codon and the region immediately upstream of it; (ii) a DNA fragment encompassing the *gfp* gene, the origin of replication, the kanamycin resistance cassette, and a 700 bp region immediately downstream of the *citZ* stop codon from pJLG341, amplified with primers oJLG95 and oER1316.

**pER394**. This plasmid was constructed by assembling the following four fragments by Gibson Assembly (New England Biolabs): (i) 3' region of *leuA* coding sequence (not including the stop codon) amplified with primers oER1341 and oER1342 from genomic DNA of *B. subtilis* PY79; (ii) *gfpΩkan* fragment amplified with primers oJLG7 and oJLG77 from pJLG38 (Shin et al. 2015); (iii) region immediately downstream of the *leuA* stop codon, amplified with primers oER1343 and oER1344 from genomic DNA of *B. subtilis* PY79; and (iv) a DNA fragment encompassing the spectinomycin resistance gene, the origin of replication, and the ampicillin resistance gene from pDG1662 (Guérout-Fleury et al. 1996), amplified with primers oJLG95 and oJLG96.

**pER397**. This plasmid was constructed by assembling the following four fragments by Gibson Assembly (New England Biolabs): (i) 3' region of *carB* coding sequence (not including the stop codon) amplified with primers oER1383 and oER1384 from genomic DNA of *B. subtilis* PY79; (ii) *gfpΩkan* fragment amplified with primers oJLG7 and oJLG77 from pJLG38 (Shin et al. 2015); (iii) region immediately downstream of the *carB* stop codon, amplified with primers oER1385 and oER1386 from genomic DNA of *B. subtilis* PY79; and (iv) a DNA fragment encompassing the spectinomycin resistance gene, the origin of replication, and the ampicillin resistance gene from pDG1662 (Guérout-Fleury et al. 1996), amplified with primers oJLG95 and oJLG96.

**pER398**. This plasmid was constructed by assembling the following four fragments by Gibson Assembly (New England Biolabs): (i) 3' region of *ilvB* coding sequence (not including the stop codon) amplified with primers oER1389 and oER1390 from genomic DNA of *B. subtilis* PY79; (ii) *gfpΩkan* fragment amplified with primers oJLG7 and oJLG77 from pJLG38 (Shin et al. 2015); (iii) region immediately downstream of the *ilvB* stop codon, amplified with primers oER1391 and oER1392 from genomic DNA of *B. subtilis* PY79; and (iv) a DNA fragment encompassing the spectinomycin resistance gene, the origin of replication, and the ampicillin resistance gene from pDG1662 (Guérout-Fleury et al. 1996), amplified with primers oJLG95 and oJLG96.

**pER403**. This plasmid was constructed by assembling the following four fragments by Gibson Assembly (New England Biolabs): (i) 3' region of *pckA* coding sequence (not including the stop codon) amplified with primers oER1401 and oER1405 from genomic DNA of *B. subtilis* PY79; (ii) *gfpΩkan* fragment amplified with primers oJLG7 and oJLG77 from pJLG38 (Shin et al. 2015); (iii) region immediately downstream of the *pckA* stop codon, amplified with primers oER1403 and oER1404 from genomic DNA of *B. subtilis* PY79; and (iv) a DNA fragment encompassing the spectinomycin resistance gene, the origin of replication, and the ampicillin resistance gene from pDG1662 (Guérout-Fleury et al. 1996), amplified with primers oJLG95 and oJLG96.

**pER406**. This plasmid was constructed by assembling the following four fragments by Gibson Assembly (New England Biolabs): (i) 3' region of *pdhD* coding sequence (not including the stop codon) amplified with primers oER1415 and oER1419 from genomic DNA of *B. subtilis* PY79; (ii) *gfpΩkan* fragment amplified with primers oJLG7 and oJLG77 from pJLG38 (Shin et al. 2015); (iii) region immediately downstream of the *pdhD* stop codon, amplified with primers oER1417 and oER1418 from genomic DNA of *B. subtilis* PY79; and (iv) a DNA fragment encompassing the spectinomycin resistance gene, the origin of replication, and the ampicillin resistance gene from pDG1662 (Guérout-Fleury et al. 1996), amplified with primers oJLG95 and oJLG96.

**pER422**. This plasmid was constructed by assembling the following four fragments by Gibson Assembly (New England Biolabs): (i) 3' region of *ndh* coding sequence (not including the stop codon) amplified with primers oER728 and oER736 from genomic DNA of *B. subtilis* PY79; (ii) *gfpΩkan* fragment amplified with primers oJLG7 and oJLG77 from pJLG38 (Shin et al. 2015); (iii) region immediately downstream of the *ndh* stop codon, amplified with primers oER735 and oER732 from genomic DNA of *B. subtilis* PY79; and (iv) a DNA fragment encompassing the spectinomycin resistance gene, the origin of replication, and the ampicillin resistance gene from pDG1662 (Guérout-Fleury et al. 1996), amplified with primers oJLG95 and oJLG96.

**pER423.** This plasmid was constructed by assembling the following four fragments by Gibson Assembly (New England Biolabs): (i) 3' region of *sigF* coding sequence (not including the stop codon) amplified with primers oJLG440 and oER1435 from genomic DNA of *B. subtilis* PY79; (ii) *gfpΩkan* fragment amplified with primers oJLG7 and oJLG77 from pJLG38 (Shin et al. 2015); (iii) region immediately downstream of the *sigF* stop codon, amplified with primers oJLG442 and oJLG443 from genomic DNA of *B. subtilis* PY79; and (iv) a DNA fragment encompassing the spectinomycin resistance gene, the origin of replication, and the ampicillin resistance gene from pDG1662 (Guérout-Fleury et al. 1996), amplified with primers oJLG95 and oJLG96.

**pER424**. This plasmid was constructed by assembling the following four fragments by Gibson Assembly (New England Biolabs): (i) 3' region of *glpX* coding sequence (not including the stop codon) amplified with primers oER1454 and oER1455 from genomic DNA of *B. subtilis* PY79; (ii) *gfpΩkan* fragment amplified with primers oJLG7 and oJLG77 from pJLG38 (Shin et al. 2015); (iii) region immediately downstream of the *glpX* stop codon, amplified with primers oER1456 and oER1457 from genomic DNA of *B. subtilis* PY79; and (iv) a DNA fragment encompassing the spectinomycin resistance gene, the origin of replication, and the ampicillin resistance gene from pDG1662 (Guérout-Fleury et al. 1996), amplified with primers oJLG95 and oJLG96.

**pER425**. This plasmid was constructed by assembling the following four fragments by Gibson Assembly (New England Biolabs): (i) 3' region of *acoC* coding sequence (not including the stop codon) amplified with primers oER1460 and oER1461 from genomic DNA of *B. subtilis* PY79; (ii) *gfpΩkan* fragment amplified with primers oJLG7 and oJLG77 from pJLG38 (Shin et al. 2015); (iii) region immediately downstream of the *acoC* stop codon, amplified with primers oER1462 and oER1463 from genomic DNA of *B. subtilis* PY79; and (iv) a DNA fragment encompassing the spectinomycin resistance gene, the origin of replication, and the ampicillin resistance gene from pDG1662 (Guérout-Fleury et al. 1996), amplified with primers oJLG95 and oJLG96.

**pER426**. This plasmid was constructed by assembling the following four fragments by Gibson Assembly (New England Biolabs): (i) 3' region of *lutC* coding sequence (not including the stop codon) amplified with primers oER1466 and oER1467 from genomic DNA of *B. subtilis* PY79; (ii) *gfpΩkan* fragment amplified with primers oJLG7 and oJLG77 from pJLG38 (Shin et al. 2015); (iii) region immediately downstream of the *lutC* stop codon, amplified with primers oER1468 and oER1469 from genomic DNA of *B. subtilis* PY79; and (iv) a DNA fragment encompassing the spectinomycin resistance gene, the origin of replication, and the ampicillin resistance gene from pDG1662 (Guérout-Fleury et al. 1996), amplified with primers oJLG95 and oJLG96.

**pER427**. This plasmid was constructed by assembling the following four fragments by Gibson Assembly (New England Biolabs): (i) 3' region of *acsA* coding sequence (not including the stop codon) amplified with primers oER1472 and oER1473 from genomic DNA of *B. subtilis* PY79; (ii) *gfpΩkan* fragment amplified with primers oJLG7 and oJLG77 from pJLG38 (Shin et al. 2015); (iii) region immediately downstream of the *acsA* stop codon, amplified with primers oER1474 and oER1475 from genomic DNA of *B. subtilis* PY79; and (iv) a DNA fragment encompassing the spectinomycin resistance gene, the origin of replication, and the ampicillin resistance gene from pDG1662 (Guérout-Fleury et al. 1996), amplified with primers oJLG95 and oJLG96.

**pER430**. This plasmid was constructed by assembling the following four fragments by Gibson Assembly (New England Biolabs): (i) 3' region of *pyrAA* coding sequence (not including the stop codon) amplified with primers oER1496 and oER1497 from genomic DNA of *B. subtilis* PY79; (ii) *gfpΩkan* fragment amplified with primers oJLG7 and oJLG77 from pJLG38 (Shin et al. 2015); (iii) region immediately downstream of the *pyrAA* stop codon, amplified with primers oER1498 and oER1499 from genomic DNA of *B. subtilis* PY79; and (iv) a DNA fragment encompassing the spectinomycin resistance gene, the origin of replication, and the ampicillin resistance gene from pDG1662 (Guérout-Fleury et al. 1996), amplified with primers oJLG95 and oJLG96.

**pER431**. This plasmid was constructed by assembling the following four fragments by Gibson Assembly (New England Biolabs): (i) 3' region of *pyrAB* coding sequence (not including the stop codon) amplified with primers oER1502 and oER1503 from genomic DNA of *B. subtilis* PY79; (ii) *gfpΩkan* fragment amplified with primers oJLG7 and oJLG77 from pJLG38 (Shin et al. 2015); (iii) region immediately downstream of the *pyrAB* stop codon, amplified with primers oER1504 and oER1505 from genomic DNA of *B. subtilis* PY79; and (iv) a DNA fragment encompassing the spectinomycin resistance gene, the origin of replication, and the ampicillin resistance gene from pDG1662 (Guérout-Fleury et al. 1996), amplified with primers oJLG95 and oJLG96.

**pER435.** This plasmid was constructed by mutagenic PCR using a QuickChange primer oER1520 on pER315.

**pER437.** This plasmid was constructed by assembling the following 2 fragments by Gibson Assembly (New England Biolabs): (i) a DNA fragment encompassing the spectinomycin resistance gene, the origin of replication, and the ampicillin resistance gene from pDG1662 and the *yjbA* downstream homology region oJLG96 and oER1523 on plasmid pER344; (ii) *yjbA(E116K)Ωerm* amplified using primers oER1522 and oER1526 on genomic DNA of BER1859.

**pER441**. This plasmid was constructed by assembling the following four fragments by Gibson Assembly (New England Biolabs): (i) 3' region of *ybgE* coding sequence (not including the stop codon) amplified with primers oER1551 and oER1552 from genomic DNA of *B. subtilis* PY79; (ii) *gfpΩkan* fragment amplified with primers oJLG7 and oJLG77 from pJLG38 (Shin et al. 2015); (iii) region immediately downstream of the *ybgE* stop codon, amplified with primers oER1553 and oER1554 from genomic DNA of *B. subtilis* PY79; and (iv) a DNA fragment encompassing the spectinomycin resistance gene, the origin of replication, and the ampicillin resistance gene from pDG1662 (Guérout-Fleury et al. 1996), amplified with primers oJLG95 and oJLG96.

**pER442**. This plasmid was constructed by assembling the following four fragments by Gibson Assembly (New England Biolabs): (i) 3' region of *argJ* coding sequence (not including the stop codon) amplified with primers oER1533 and oER1534 from genomic DNA of *B. subtilis* PY79; (ii) *gfpΩkan* fragment amplified with primers oJLG7 and oJLG77 from pJLG38 (Shin et al. 2015); (iii) region immediately downstream of the *argJ* stop codon, amplified with primers oER1535 and oER1536 from genomic DNA of *B. subtilis* PY79; and (iv) a DNA fragment encompassing the spectinomycin resistance gene, the origin of replication, and the ampicillin resistance gene from pDG1662 (Guérout-Fleury et al. 1996), amplified with primers oJLG95 and oJLG96.

**pER443**. This plasmid was constructed by assembling the following four fragments by Gibson Assembly (New England Biolabs): (i) 3' region of *argB* coding sequence (not including the stop codon) amplified with primers oER1539 and oER1540 from genomic DNA of *B. subtilis* PY79; (ii) *gfpΩkan* fragment amplified with primers oJLG7 and oJLG77 from pJLG38 (Shin et al. 2015); (iii) region immediately downstream of the *argB* stop codon, amplified with primers oER1541 and oER1542 from genomic DNA of *B. subtilis* PY79; and (iv) a DNA fragment encompassing the spectinomycin resistance gene, the origin of replication, and the ampicillin resistance gene from pDG1662 (Guérout-Fleury et al. 1996), amplified with primers oJLG95 and oJLG96.

**pER444**. This plasmid was constructed by assembling the following four fragments by Gibson Assembly (New England Biolabs): (i) 3' region of *argD* coding sequence (not including the stop codon) amplified with primers oER1557 and oER1558 from genomic DNA of *B. subtilis* PY79; (ii) *gfpΩkan* fragment amplified with primers oJLG7 and oJLG77 from pJLG38 (Shin et al. 2015); (iii) region immediately downstream of the *argD* stop codon, amplified with primers oER1559 and oER1560 from genomic DNA of *B. subtilis* PY79; and (iv) a DNA fragment encompassing the spectinomycin resistance gene, the origin of replication, and the ampicillin resistance gene from pDG1662 (Guérout-Fleury et al. 1996), amplified with primers oJLG95 and oJLG96.

**pER447**. This plasmid was constructed by assembling the following four fragments by Gibson Assembly (New England Biolabs): (i) 3' region of *yxbC* coding sequence (not including the stop codon) amplified with primers oER1565 and oER1566 from genomic DNA of *B. subtilis* PY79; (ii) *gfpΩkan* fragment amplified with primers oJLG7 and oJLG77 from pJLG38 (Shin et al. 2015); (iii) region immediately downstream of the *yxbC* stop codon, amplified with primers oER1567 and oER1568 from genomic DNA of *B. subtilis* PY79; and (iv) a DNA fragment encompassing the spectinomycin resistance gene, the origin of replication, and the ampicillin resistance gene from pDG1662 (Guérout-Fleury et al. 1996), amplified with primers oJLG95 and oJLG96.

**pER452**. This plasmid was constructed by assembling the following four fragments by Gibson Assembly (New England Biolabs): (i) 3' region of *purL* coding sequence (not including the stop codon) amplified with primers oER1589 and oER1590 from genomic DNA of *B. subtilis* PY79; (ii) *gfpΩkan* fragment amplified with primers oJLG7 and oJLG77 from pJLG38 (Shin et al. 2015); (iii) region immediately downstream of the *purL* stop codon, amplified with primers oER1591 and oER1592 from genomic DNA of *B. subtilis* PY79; and (iv) a DNA fragment encompassing the spectinomycin resistance gene, the origin of replication, and the ampicillin resistance gene from pDG1662 (Guérout-Fleury et al. 1996), amplified with primers oJLG95 and oJLG96.

**pER454.** This plasmid was constructed by assembling the following four fragments by Gibson Assembly (New England Biolabs): (i) 3' region of *divIVA* coding sequence (not including the stop codon) amplified with primers oER1607 and oER1608 from genomic DNA of *B. subtilis* PY79; (ii) *gfpΩkan* fragment amplified with primers oJLG7 and oJLG77 from pJLG38 (Shin et al. 2015); (iii) region immediately downstream of the *divIVA* stop codon, amplified with primers oER1609 and oER1610 from genomic DNA of *B. subtilis* PY79; and (iv) a DNA fragment encompassing the spectinomycin resistance gene, the origin of replication, and the ampicillin resistance gene from pDG1662 (Guérout-Fleury et al. 1996), amplified with primers oJLG95 and oJLG96.

**pER455**. This plasmid was constructed by assembling the following four fragments by Gibson Assembly (New England Biolabs): (i) 3' region of *thrC* coding sequence (not including the stop codon) amplified with primers oER1613 and oER1614 from genomic DNA of *B. subtilis* PY79; (ii) *gfpΩkan* fragment amplified with primers oJLG7 and oJLG77 from pJLG38 (Shin et al. 2015); (iii) region immediately downstream of the *thrC* stop codon, amplified with primers oER1615 and oER1616 from genomic DNA of *B. subtilis* PY79; and (iv) a DNA fragment encompassing the spectinomycin resistance gene, the origin of replication, and the ampicillin resistance gene from pDG1662 (Guérout-Fleury et al. 1996), amplified with primers oJLG95 and oJLG96.

**pER456**. This plasmid was constructed by assembling the following four fragments by Gibson Assembly (New England Biolabs): (i) 3' region of *hpf* coding sequence (not including the stop codon) amplified with primers oER1619 and oER1620 from genomic DNA of *B. subtilis* PY79; (ii) *gfpΩkan* fragment amplified with primers oJLG7 and oJLG77 from pJLG38 (Shin et al. 2015); (iii) region immediately downstream of the *hpf* stop codon, amplified with primers oER1621 and oER1622 from genomic DNA of *B. subtilis* PY79; and (iv) a DNA fragment encompassing the spectinomycin resistance gene, the origin of replication, and the ampicillin resistance gene from pDG1662 (Guérout-Fleury et al. 1996), amplified with primers oJLG95 and oJLG96.

**pER458.** This plasmid was constructed by assembling the following four fragments by Gibson Assembly (New England Biolabs): (i) a *ccpA* upstream homology fragment amplified with primers oER1631 and oER1632 from genomic DNA of *B. subtilis* PY79; (ii) a erythromycin resistance gene amplified with primers oER419 and oER420 from BER0389; (iii) a *ccpA* downstream homology fragment amplified with primers oER1633 and oER1517 from genomic DNA of *B. subtilis* PY79; and (iv) a DNA fragment encompassing the spectinomycin resistance gene, the origin of replication, and the ampicillin resistance gene from pDG1662 (Guérout-Fleury et al. 1996), amplified with primers oJLG95 and oJLG96.

**pER461**. This plasmid was constructed by assembling the following four fragments by Gibson Assembly (New England Biolabs): (i) 3' region of *metC* coding sequence (not including the stop codon) amplified with primers oER1659 and oER1660 from genomic DNA of *B. subtilis* PY79; (ii) *gfpΩkan* fragment amplified with primers oJLG7 and oJLG77 from pJLG38 (Shin et al. 2015); (iii) region immediately downstream of the *metC* stop codon, amplified with primers oER1661 and oER1662 from genomic DNA of *B. subtilis* PY79; and (iv) a DNA fragment encompassing the spectinomycin resistance gene, the origin of replication, and the ampicillin resistance gene from pDG1662 (Guérout-Fleury et al. 1996), amplified with primers oJLG95 and oJLG96.

**pER462**. This plasmid was constructed by assembling the following four fragments by Gibson Assembly (New England Biolabs): (i) 3' region of *cysC* coding sequence (not including the stop codon) amplified with primers oER1665 and oER1666 from genomic DNA of *B. subtilis* PY79; (ii) *gfpΩkan* fragment amplified with primers oJLG7 and oJLG77 from pJLG38 (Shin et al. 2015); (iii) region immediately downstream of the *cysC* stop codon, amplified with primers oER1667 and oER1668 from genomic DNA of *B. subtilis* PY79; and (iv) a DNA fragment encompassing the spectinomycin resistance gene, the origin of replication, and the ampicillin resistance gene from pDG1662 (Guérout-Fleury et al. 1996), amplified with primers oJLG95 and oJLG96.

**pER463**. This plasmid was constructed by assembling the following four fragments by Gibson Assembly (New England Biolabs): (i) 3' region of *sat* coding sequence (not including the stop codon) amplified with primers oER1671 and oER1672 from genomic DNA of *B. subtilis* PY79; (ii) *gfpΩkan* fragment amplified with primers oJLG7 and oJLG77 from pJLG38 (Shin et al. 2015); (iii) region immediately downstream of the *sat* stop codon, amplified with primers oER1673 and oER1674 from genomic DNA of *B. subtilis* PY79; and (iv) a DNA fragment encompassing the spectinomycin resistance gene, the origin of replication, and the ampicillin resistance gene from pDG1662 (Guérout-Fleury et al. 1996), amplified with primers oJLG95 and oJLG96.

**pER464.** This plasmid was constructed by assembling the following 2 fragments by Gibson Assembly (New England Biolabs): (i) a 5706 bp fragment of *lacA::PspoIIEΩerm* amplified from pER128 (Riley et al 2018) using primers oER1104 and oER1105; (ii) *citZ-gfpΩkan* fragment amplified with primers oER1656 and oER1657 from genomic DNA of JLG2113 (Riley et al. 2021).

**pER465.** This plasmid was constructed by assembling the following fragment by Gibson Assembly (New England Biolabs): xylA::gfp*Ωcat* reverse amplified using primers oER1636 and oER1658 from plasmid pER360.

**pER467.** This plasmid was constructed by assembling the following 2 fragments by Gibson Assembly (New England Biolabs): (i) xylA::gfp*Ωcat* reverse amplified using primers oER1636 and oER1658 from plasmid pER360; (ii) *citZ-gfp* fragment amplified with primers oER1656 and oER1657 from genomic DNA of JLG2113 (Shin et al. 2015).

**pJAL011.** This plasmid was constructed by assembling the following four fragments by Gibson Assembly (New England Biolabs): (i) 3' region of *etfA* coding sequence (not including the stop codon) amplified with primers oJAL007 and oJAL008 from genomic DNA of *B. subtilis* PY79; (ii) *gfpΩkan* fragment amplified with primers oJLG7 and oJLG77 from pJLG38 (Shin et al. 2015); (iii) region immediately downstream of the *etfA* stop codon, amplified with primers oJAL009 and oJAL010 from genomic DNA of *B. subtilis* PY79; and (iv) a DNA fragment encompassing the spectinomycin resistance gene, the origin of replication, and the ampicillin resistance gene from pDG1662 (Guérout-Fleury et al. 1996), amplified with primers oJLG95 and oJLG96.

**pJLG396**. Constructed by assembling the following fragments by Gibson assembly: pER226 (Lopez-Garrido et al. 2018) inverse PCR fragment amplified with primers oJLG1406 and oJLG1407, and *erm* cassette, amplified with oJLG1408 and oJLG1409 from pDG1731 (Guérout-Fleury et al. 1996).

**pJLG1214**. This plasmid was constructed by assembling the following four fragments by Gibson isothermal assembly (New England Biolabs): (i) 3' region of the *dat* coding sequence, not including the stop codon, amplified from genomic DNA of *B. subtilis* PY79 with primers oJLG3036 and oJLG3037; (ii) sfGFP-loxP-Kan-loxP fragment amplified from pJLG396 with primers oJLG7 and oJLG1699; (iii) region immediately downstream of dat coding sequence, amplified from genomic DNA of *B. subtilis* PY79 with primers oJLG3038 and oJLG3039; and (iv) a DNA fragment encompassing the spectinomycin resistant gene, the origin of replication, and the beta-lactamase gene from pDG1662, amplified with primers oJLG1696 and oJLG96.

*Details of proteomic sample preparation*
MS samples were prepared from the spore samples and analyzed by the University of California, San Diego Biomolecular and Proteomics MS Facility (http://bpmsf.ucsd.edu/).

Protein digestion Protocol: The spore pellet was resuspended in 600ul of 8 M Urea in 100mM Tris pH 8.0 by vortexing for 5-10 minutes. TCEP was added to final concentration of 10 mM. The sample was then frozen overnight at -20C to help solubilize the proteins. The solution was thawed and vortexed for another 5 minutes until the solution became clear. Chloro-acetamide solution was added to final concentration of 40 mM and vortexed for 5 minutes. An equal volume of 50mM Tris pH 8.0 was added to the sample to reduce the urea concentration to 4M. It is ideal to carry out a protein concentration measurement at this step in preparation of protease addition. LysC was added in 1:500 ratio of LysC to protein content and the samples were incubated at 37C in a rotating incubator for 4-6 hr. An equal volume of 50mM Tris pH 8.0 was added to the sample to reduce the urea concentration to 2 M. Trypsin was in 1:50 ratio of trypsin to protein content. Next day the solution was acidified using TFA (0.5% TFA final concentration) and vortexed for 5 minutes. The sample was centrifuged at 15 700g for 5 min to obtain aqueous and organic phases. The lower aqueous phase was collected and desalted using 100 mg C18-StageTips as described by the manufacturer protocol. The peptide concentration of sample was measured using BCA after resuspension iTRAQ dissolution buffer.

iTRAQ labeling: 2 tags of Thermo Scientific™ TMT™ 10plex (Catalog number: 90110) were used for sample labeling as described by manufacturer.

| | |
|---|---|
| C1 | • TMT10-129C |
| E1 | • TMT10-130N |

High pH fractionation: Pierce™ High pH Reversed-Phase Peptide Fractionation Kit (Pierce™ High pH Reversed-Phase Peptide Fractionation Kit Catalog number: 84868) was used. Fractionation protocol as described by the manufacturer.
LC-MS-MS: Each fraction was analyzed by ultra-high-pressure liquid chromatography (UPLC) coupled with tandem mass spectroscopy (LC-MS/MS) using nano-spray ionization. The nano-spray ionization experiments were performed using an Orbitrap fusion Lumos hybrid mass spectrometer (Thermo) interfaced with nano-scale reversed-phase UPLC (Thermo Dionex UltiMate™ 3000 RSLC nano System) using a 25 cm, 75-micron ID glass capillary packed with 1.7-µm C18 (130) BEH™ beads (Waters corporation). Peptides were eluted from the C18 column into the mass spectrometer using a linear gradient (5–80%) of ACN (Acetonitrile) at a flow rate of 375 µl/min for 120 min. The buffers used to create the ACN gradient were: Buffer A (98% $H_2O$, 2% ACN, 0.1% formic acid) and Buffer B (100% ACN, 0.1% formic acid). Mass spectrometer parameters are as follows; an MS1 survey scan using the orbitrap detector (mass range (m/z): 400-1500 (using quadrupole isolation), 60000 resolution setting, spray voltage of 2200 V, Ion transfer tube temperature of 275 C, AGC target of 400000, and maximum injection time of 50 ms) was followed by data dependent scans (top speed for most intense ions), with charge state set to only include +2-5 ions, and 5 second exclusion time, while selecting ions with minimal intensities of 50000 at in which the collision event was carried out in the high energy collision cell (HCD Collision Energy of 38%) and the first quadrupole isolation window was set at 0.7 (m/z). The fragment masses were analyzed in the Orbi-trap mass analyzer with mass resolution setting of 15000 (With ion trap scan rate of turbo, first mass m/z was 100, AGC Target 20000 and maximum

injection time of 22ms). Protein identification and quantification was carried out using Peaks Studio 8.5 (Bioinformatics solutions Inc.).
